# Low-cost, bottom-up fabrication of large-scale single-molecule nanoarrays by DNA origami placement

**DOI:** 10.1101/2020.08.14.250951

**Authors:** Rishabh M. Shetty, Sarah R. Brady, Paul W. K. Rothemund, Rizal F. Hariadi, Ashwin Gopinath

## Abstract

Large-scale nanoarrays of single biomolecules enable high-throughput assays while unmasking the underlying heterogeneity within ensemble populations. Until recently, creating such grids which combine the unique advantages of microarrays and single-molecule experiments (SMEs) has been particularly challenging due to the mismatch between the size of these molecules and the resolution of top-down fabrication techniques. DNA Origami Placement (DOP) combines two powerful techniques to address this issue: (*i*) DNA origami, which provides a ∼ 100-nm self-assembled template for single-molecule organization with 5 nm resolution, and (*ii*) top-down lithography, which patterns these DNA nanostructures, transforming them into functional nanodevices *via* large-scale integration with arbitrary substrates. Presently, this technique relies on state-of-the-art infrastructure and highly-trained personnel, making it prohibitively expensive for researchers. Here, we introduce a bench-top technique to create meso-to-macro-scale DNA origami nanoarrays using self-assembled colloidal nanoparticles, thereby circumventing the need for top-down fabrication. We report a maximum yield of 74%, two-fold higher than the statistical limit of 37% imposed on non-specific molecular loading alternatives. Furthermore, we provide a proof-of-principle for the ability of this nanoarray platform to transform traditionally low-throughput, stochastic, single-molecule assays into high-throughput, deterministic ones, without compromising data quality. Our approach has the potential to democratize single-molecule nanoarrays and demonstrates their utility as a tool for biophysical assays and diagnostics.

## Introduction

Bulk measurements yield little information about the heterogeneity prevalent at the single-molecule level^1,2^. The interest in gaining quantitative and mechanistic insight into these molecular processes spurred the development of novel biophysical and analytical single-molecule methods over the past few decades^2–4^. Since the introduction of Total Internal Reflection Fluorescence (TIRF) microscopy^5,6^, single-molecule experiments of biomolecular kinetics, conformational fluctuations, and folding mechanisms have become commonplace in biophysics laboratories^7,8^. Classical single-molecule experiments such as these are stochastic in nature^7–14^, with biophysicists lacking the ability to control where individual molecules bind on surfaces. This leads to the possibility that two or more molecules may occupy the same diffraction-limited spot, often leading to confounding data (**Fig. 1A** and **B**). Reducing the concentration of molecules to overcome this issue has the caveat of lowering experimental throughput. This concentration versus throughput conundrum is a major limitation of conventional single-molecule studies^1^. Maximizing throughput while controlling the positions of molecules-of-interest for optimal data quality on a substrate would require close-packing (**Fig. 1C**), ideally at the diffraction limit of light ≃ λ/2NA, where λ is the wavelength of excitation light, and NA is the numerical aperture of the objective lens. However, any deterministic positioning of molecules requires precise positional control on experimentally relevant substrates, and the size of most single molecules is well below the resolution of current micro-to-nanomanipulation techniques.

**Fig. 1.**
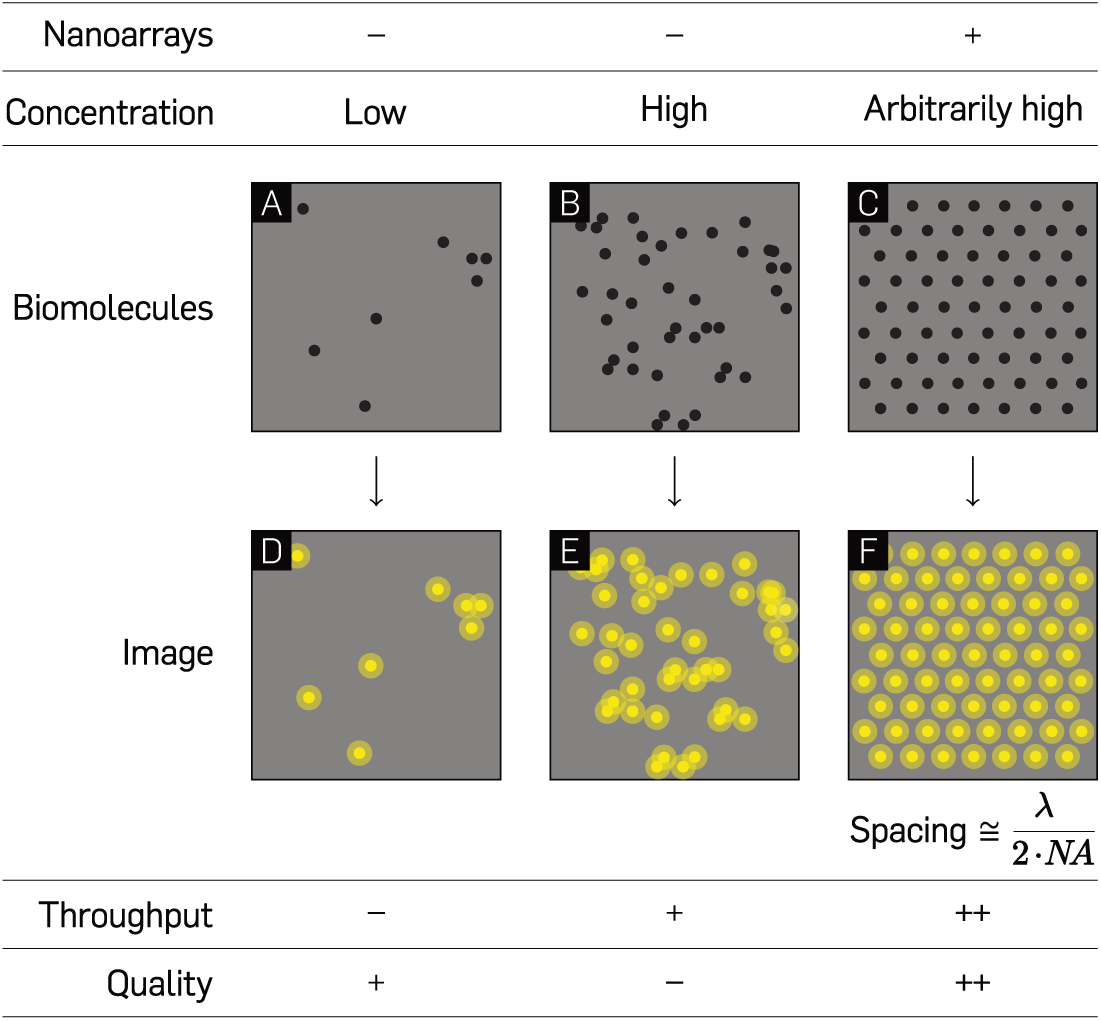
Comparison between experiments with unpatterned molecules and single-molecule nanoarrays. Hypothetical microscopy samples of low (**A** and **D**) and high (**B** and **E**) concentrations of randomly immobilized single molecules on optical substrates and their corresponding images. (**C** and **F**) Single molecule arrays on a patterned glass substrate at an inter-molecular distance marginally larger than the diffraction limit of light microscopy.

DNA origami^15^ is regarded as a molecular breadboard and bridge between the bottom-up worlds of biochemistry and the top-down world of lithography^16^. DNA origami nanotechnology is modular and spatially-programmable^17–22^; an assembled origami unit being capable of carrying up to 200 individually addressable molecules-of-interest^23–26^. In the last decade, origami nanostructures have been utilized for a myriad of applications ranging from electronic–^27,28^ and optical–devices^14,26,29,30^, to single-molecule biophysics^9–11,31,32^, biosensing^33–35^, and nanofabrication^36–40^. Being synthesized in solution, spatial stochasticity is intrinsically linked with the deposition of planar origami and their payload on glass substrates for optical experiments. A 2D DNA origami nanostructure (∼ 100 nm), however, is more than an order of magnitude larger than other molecules, which makes it amenable to lithographic manipulation and deterministic positioning.

Electron Beam Lithography-based DNA Origami Placement (DOP)^36–38^ leverages the ability of origami nanostructures – through their electrostatic or covalent coupling to mica, glass, silicon, and silicon nitride – to interface biomolecular functional moieties with the outside world for visual probing. A recent application of this method demonstrates the large-scale integration of functionalized DNA origami through placement on ∼ 100-nm binding sites with >90% single-binding efficiency for hybrid nanodevice fabrication^36^. Such a composite nano-to-micro-manipulation technique enables bi-level control— first, through the arbitrary decoration of molecules with a resolution of 5 nm on origami nanostructures, and second, by positioning the origami themselves on lithographically-patterned sites on a desired substrate. The major drawback of lithographic techniques for origami placement is their high-cost owing to the manufacturing complexity of top-down fabrication. The wide-scale utilization of such processes is therefore impractical for scientific research such as biophysics, which traditionally does not use sophisticated top-down nanofabrication.

Bottom-up, self-assembly based approaches have the unique potential to provide a framework for parallel fabrication of structures from components either too diminutive or innumerable to be handled robotically^41^. Such processes were predicted to be a cornerstone of the field of nanotechnology during its nascent stages^42^. Self-assembly techniques like nanosphere lithography (NSL), while limited in terms of their ability to create arbitrary shapes, offer a variety of advantages – they are cheap, facilitate fast, parallel-processing, and a variety of crystallization techniques exist for covering arbitrarily large surface topologies^43,44^. In NSL, a flat, hydrophilic substrate is coated with a monodisperse colloidal suspension of spheres, and upon drying, a hexagonal-close-packed(HCP) layer called a Colloidal Crystal Mask is formed. Attractive capillary forces and convective nanosphere transport are the dominant factors in the self-assembly process^43^. The order and quality of the assembled arrays are substantially affected by the rates of solvent evaporation^45,46^. Control over the temperature and the humidity of the system on a slightly tilted substrate can yield colloidal monolayers^47^. Methods such as spin-coating^48^, Langmuir-Blodgett deposition^49^, and controlled evaporation^50^ have all been used to assemble large-scale monolayers of colloidal suspensions.

Here, we present the application of NSL to the controlled placement of DNA origami nanostructures on glass substrates as a framework for the fabrication of large-scale single molecule nanoarrays. This novel method for bench-top, cleanroom-free, DNA origami placement in meso-to-macro-scale grids utilizes tunable colloidal nanosphere masks^44,51–54^ and surface chemistry. This technique is similar to previous work^55^ which patterned gold nanoparticle arrays, but here we place the emphasis on maximizing single-molecule occupancy. Another recently introduced technique of DNA origami adsorption in nanohole arrays^39^ formed using NSL performed critical process steps circuitously in a cleanroom environment and was limited to approximately 50% single occupancy with extremely long incubation periods. In the study reported here, we first establish the optimal binding site diameter for circular origami and subsequently characterize the single origami binding. We report a maximum efficiency of 74%, two-fold higher than the Poisson limit of 37% achievable with conventional, stochastic loading of single molecules^56,57^. We provide evidence for the utility of our technique by demonstrating data quality comparable with classical, stochastic super-resolution DNA-Points Accumulation for Imaging in Nanoscale Topography (DNA-PAINT)^9–11,13,31^ experiments, but with up to an order of magnitude higher throughput. This self-assembly based technique enables the highest 2-D packing efficiency, and approaches the single-molecule binding yield of top-down electron-beam lithography (EBL)-based patterning at ∼ 50X lesser cost and significantly lower complexity. It has the potential to address the concentration vs. throughput conundrum in SMEs (**Fig. 1**), and function as a robust platform for deterministic, high-throughput biophysical studies, thereby making DOP more feasible and accessible to the scientific community at large.

## Results and discussion

### Nanosphere lithography and surface chemistry transform resource-intensive DOP to a facile bench-top process

To achieve the highest theoretical packing density, we position DNA origami nanostructures through DOP onto a hexagonal array with a defined spacing that is marginally larger than the wavelength of visible light (**Fig. 2A**). We create binding sites for origami nanostructures through a self-assembly based NSL technique (**Fig. 2B–F**). In a typical experiment, upon drying of polystyrene nanospheres in a solvent-based aqueous solution, a close-packed crystalline layer of colloidal nanoparticles is observed on a slightly tilted (∼ 45°), 1 cm^2^ hydrophilic glass surface (**Fig. 2B and C**). Cross-sectional Scanning Electron Microscope (SEM) images reveal contact areas between individual nanospheres and the glass substrate that can be utilized as “masks” for bulk vapor-phase passivation with hexamethyldisilazane (HMDS) (**Fig. 2C**), similar to passivation in top-down DOP^37^. Subsequent nanosphere “lift-off” by sonication in water results in the creation of nanosphere-dependent binding sites in these masked areas. Finally, controlled origami placement is achieved by tuning pH, Mg^2+^ concentration, origami concentration, and incubation time^37^.

**Fig. 2.**
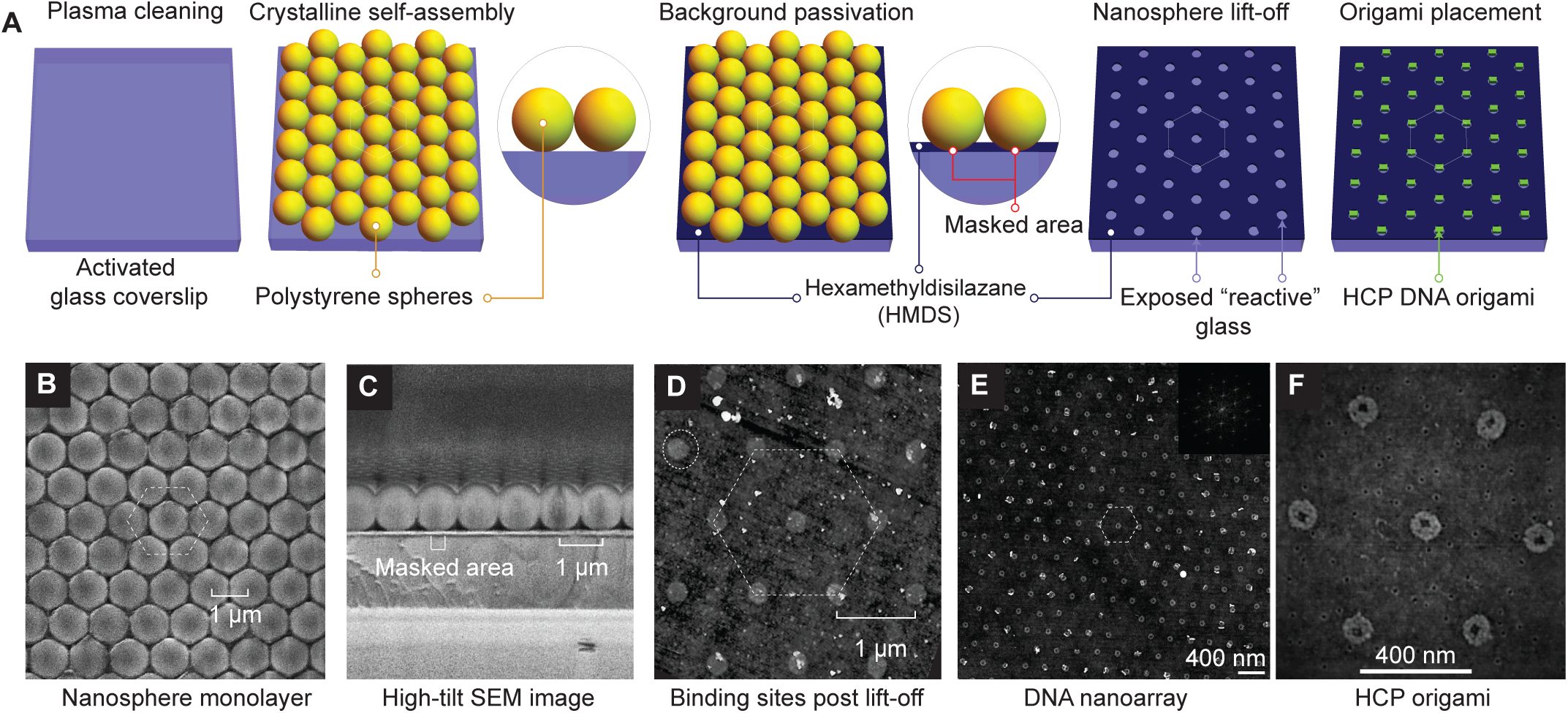
Bench-top DNA origami nanoarray fabrication. (**A**) Schematic illustration of the DNA origami patterning process through 2D nanosphere close-packing, selective passivation, lift-off, and finally, Mg^2+^-mediated origami placement. (**B, C**) SEM images of nanosphere close-packing (top view, and cross-section), respectively. (**D–F**) AFM images of binding sites (**D**), micro-scale origami placement ((**E and F**); inset: 2D FFT demonstrating close-packing). Experimental results demonstrate data analogous to schematic depiction (**A**) of process steps.

Results presented hereafter are from experiments performed at optimal values for these parameters. For the close-packed nanoparticles, closer visual inspection using (top-view) SEM images revealed continuous crystalline domains of up to 0.05 mm^2^ for 1 µm particle diameters. We found that the coverage, and uniformity of crystal domains improves with a reduction in particle diameters. Additionally, we find instances of multilayer deposition in the close-packing process. Due to the gaseous nature and minuscule size of HMDS, the additional layers of nanospheres do not affect bulk surface modification.

### Nanosphere diameter determines the spacing *s*, the binding site size *a*, and the single DNA origami occupancy

To characterize the size of the binding site as a function of nanosphere size, we imaged the binding site formation for nanospheres with diameters ranging from 200 to 1000 nm using SEM (**Fig. 3A–F**) and Atomic Force Microscopy (AFM, **Fig. 3G–L**). Within our experimental range, the spacing between neighboring binding sites *s* increases linearly with nanosphere diameter *d*_*ns*_ (*s* = 0.86*d*_*ns*_; *R*^2^ = 1.0; **Fig. 3M**). In a hexagonal-closed-packed arrangement, each nanosphere makes machanical contact with its 6 neighbors. These intermolecular interactions induce deformation and give rise to the less-than 1:1 *s/d*_*ns*_ ratio. Origami nanostructures align themselves on binding sites to maximize the number of silanol–Mg^2+^–origami bridges (**Fig. S1**), presumably by a process of 2D diffusion once they land on the surface^37^. A size match between the binding site and origami geometry is therefore paramount to maximizing single origami occupancy per binding site through steric occlusion (**Fig. 3N and O**)^37^. Each nanosphere–glass contact point deforms both bodies, with the nanosphere acting as a mask for subsequent vapor deposition of HMDS (**Fig. S2**). The nanosphere diameter *d*_*ns*_ and binding site diameter *a* relationship is defined by a Hertzian contact equation^58^ below:

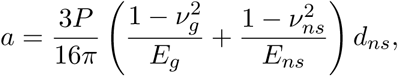

where *E*_*g*_, *E*_*ns*_ are the elastic moduli and *ν*_*g*_, *ν*_*ns*_ are the Poisson’s ratios associated with glass and nanospheres, respectively, and *P* is the applied intermolecular pressure. The binding site diameter *a* increases linearly with the nanosphere diameter *d*_*ns*_ and is given by *a* = 0.27*d*_*ns*_ (*R*^2^ = 0.99). The linear fit suggests that within the experimental range, the applied pressure *P* and the mechanical properties of the nanospheres (*ν*_*ns*_ and *E*_*ns*_) are independent of the nanosphere diameter. We find that 100-nm binding sites can be routinely fabricated using nanosphere diameters of 300–400 nm. Therefore, benchtop NSL enables controlled fabrication of arrays of binding sites for the placement of single origami molecules on glass substrates.

**Fig. 3.**
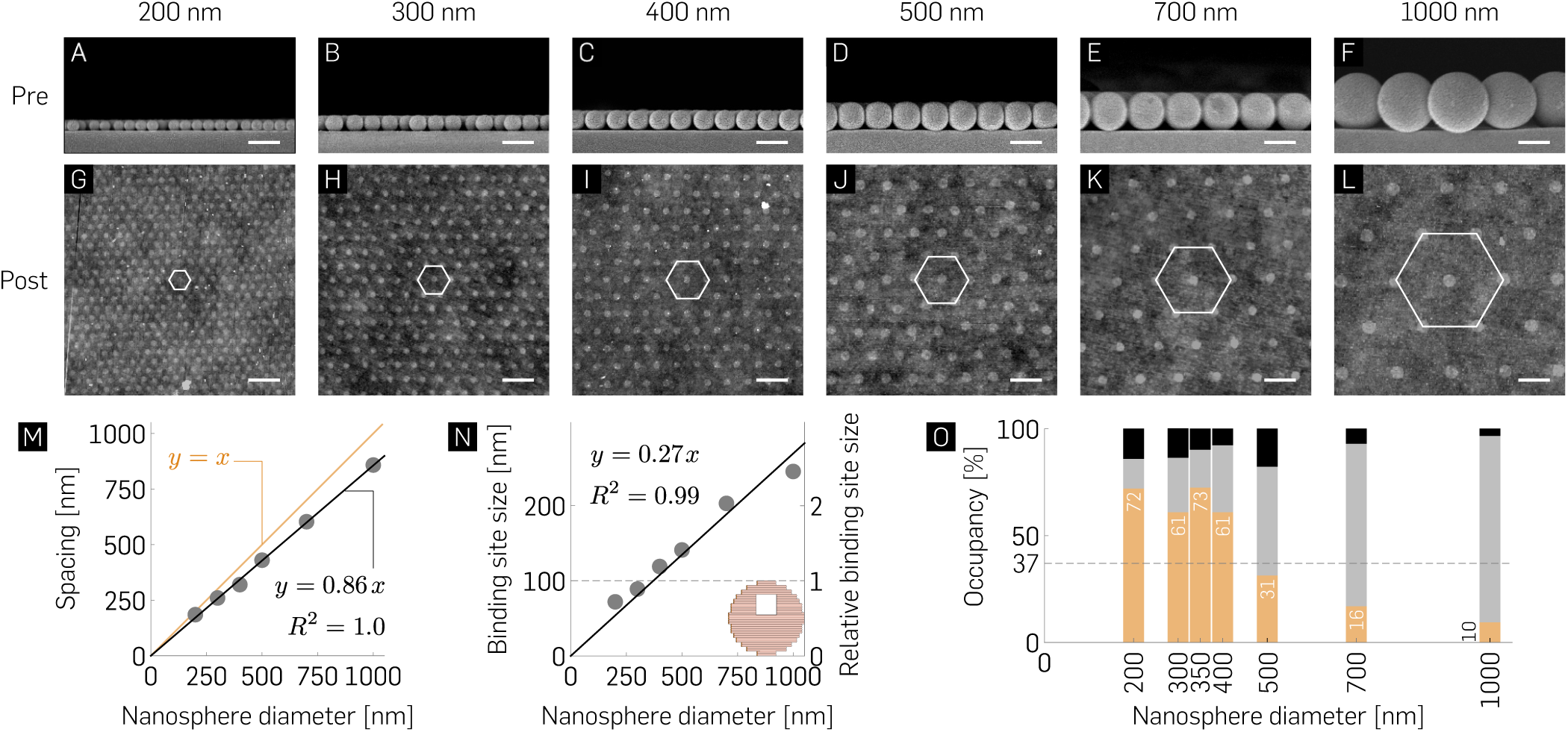
Nanosphere diameter-dependent occupancy statistics. (**A**–**L**) Cross-sectional EM images of hexagonal packing with indicated nanospheres reveal their mechanical contact with the glass surface (**A**–**F**) and their corresponding origami binding sites observed post lift-off *via* AFM imaging (**G**–**L**). Scale bars are 500 nm. (**M**,**N**) Spacing and binding site size of nanoarray patterning as a function of indicated nanosphere diameters. (**O**) Mean percentage binding of zero **(black)**, exactly one **(orange)**, and two ≥ origami **(gray)** as a function of nanosphere diameter (*N* ≥ 600) demonstrating non-Poisson statistics for single molecule binding with maximal 72.4 ± 2.14%, and 72 ± 6.84% single origami binding for 350-nm, and 200-nm nanospheres. The >70% measured probability for single origami binding is higher than the Poisson limit (horizontal dashed line, and (**Fig. S3**)). Refer to **Table S4** for the ± SD of the mean percentage bindings.

### Single origami binding statistics – ∼ 2x better than Poisson statistics, and ∼ 50X cheaper than top-down DOP

Circular origami with a square hol (**Table S2**)^59^, 100 nm across were utilized for experiments owing to their geometric similarity with binding sites. The hole served to guide the orientation of DNA origami in previously published DOP work^59^. Our experimental observations indicate a maximal, 74% single origami occupancy when the origami are 350-nm apart from each other, *i.e.* at the limit of diffraction for light microscope. Incubation conditions such as time and origami concentration were altered based on nanosphere diameter used; smaller nanospheres produce a larger number of binding sites and therefore require higher values of both these parameters. The pH (8.3–8.4) and Mg^2+^ (40 mM) concentration remained constant for all experiments reported herein. All of the micrographs presented here were obtained via imaging on an (ethanol-) dehydrated substrate (**Section S2**). While the efficiency of single-molecule occupancy reported here is lower than the >95% previously reported using EBL, we argue that the ∼ 20% occupancy difference is offset by the technique’s simplicity and ∼ 50X lower cost (**Table S3**). Results reported here corroborate our prediction that the highest single origami occupancy values would be observed around the 300–400-nm nanosphere diameter range owing to the origami and binding site geometries being almost identical to each other. Further, we find ∼ 100% occupancy of all binding sites under optimal incubation conditions. Similar to a previous study^37^, our measurement statistics likely underestimate the number of single and multiple bindings of origami on the binding sites and are, in fact, a more comprehensive reflection of the fabrication process quality. Comparable to this previous study, we found only a fractional drop in single binding efficiency in slightly undersized sites as a result of using 200 nm nanospheres. The characterization process suggests a trade-off involved in the selection of appropriate nanosphere diameters with respect to the following parameters: throughput, single origami binding efficiency, and diffraction-limited experimental observation. We explored origami placement at lower Mg^2+^ (15 mM) concentration and preliminary results (**Fig. S4**) indicate that origami binding efficiency can be optimized by rationally tuning the primary global parameters. Variability associated with placement results can be attributed, in part, to manual washing steps prior to drying and AFM characterization. An automated washing process was implemented, and preliminary experiments demonstrate 66% single origami occupancy with the automatic wash setup. The setup comprises a peristaltic pump and 3-D printed tube holder (**Fig. S5**) for positional alignment between runs without manual intervention.

### Nanoarray platform is robust at low divalent cation concentrations

Single-molecule experiments for studying dynamic events are generally performed at less than 10 mM divalent cation concentrations that are physiologically relevant and minimize the formation of biomolecular aggregates. On the contrary, DNA origami nanostructures are traditionally synthesized and stored in *>*10 mM Mg^2+^ buffers for electrostatic screening. Therefore, we explored the robustness of the nanoarray platform to assess its relevance for biophysical experiments conducted at low divalent cation concentrations. We first performed control experiments on multiple glass substrates to ascertain the quality of random immobilization of circular origami (250 pM, 30-min incubation) suspended in varying Mg^2+^ concentrations (1–40 mM) and observed that a minimum of 5 mM Mg^2+^ was required to stabilize origami on an activated glass surface post-rehydration (**Fig. 4A–C**). Next, we randomly deposited origami suspended in placement buffer (40 mM Mg^2+^, pH 8.3), (ethanol-) dehydrated the surface for AFM characterization, rehydrated the origami in buffer with 1 mM-40 mM Mg^2+^ for two hours, dehydrated once more for imaging, and observed little-to-no apparent change in the quality of origami immobilization under AFM (**Fig. 4D–F**). Finally, to confirm that this process translated effectively to programmatic placement, we patterned origami on a 700-nm “grid” (40 mM Mg^2+^, pH 8.3, 300 pM), dehydrated the surface, rehydrated for two hours in 1–40mM Mg^2+^ concentrations, and dehydrated in ethanol once more. We observed high-quality grids via fluorescence micrographs as assessed by their 2D-Fourier Transforms (**Fig. 4G–I**) which confirmed the conservation of spatial conformation over time. These results demonstrate the robustness of this platform at low salt concentrations and validate its use for physiologically relevant single-molecule biophysics experiments. Prior to each dehydration step, the substrate was washed for one minute in 1x TAE (12.5 mM Mg^2+^) buffer to remove any non-specifically bound origami.

**Fig. 4.**
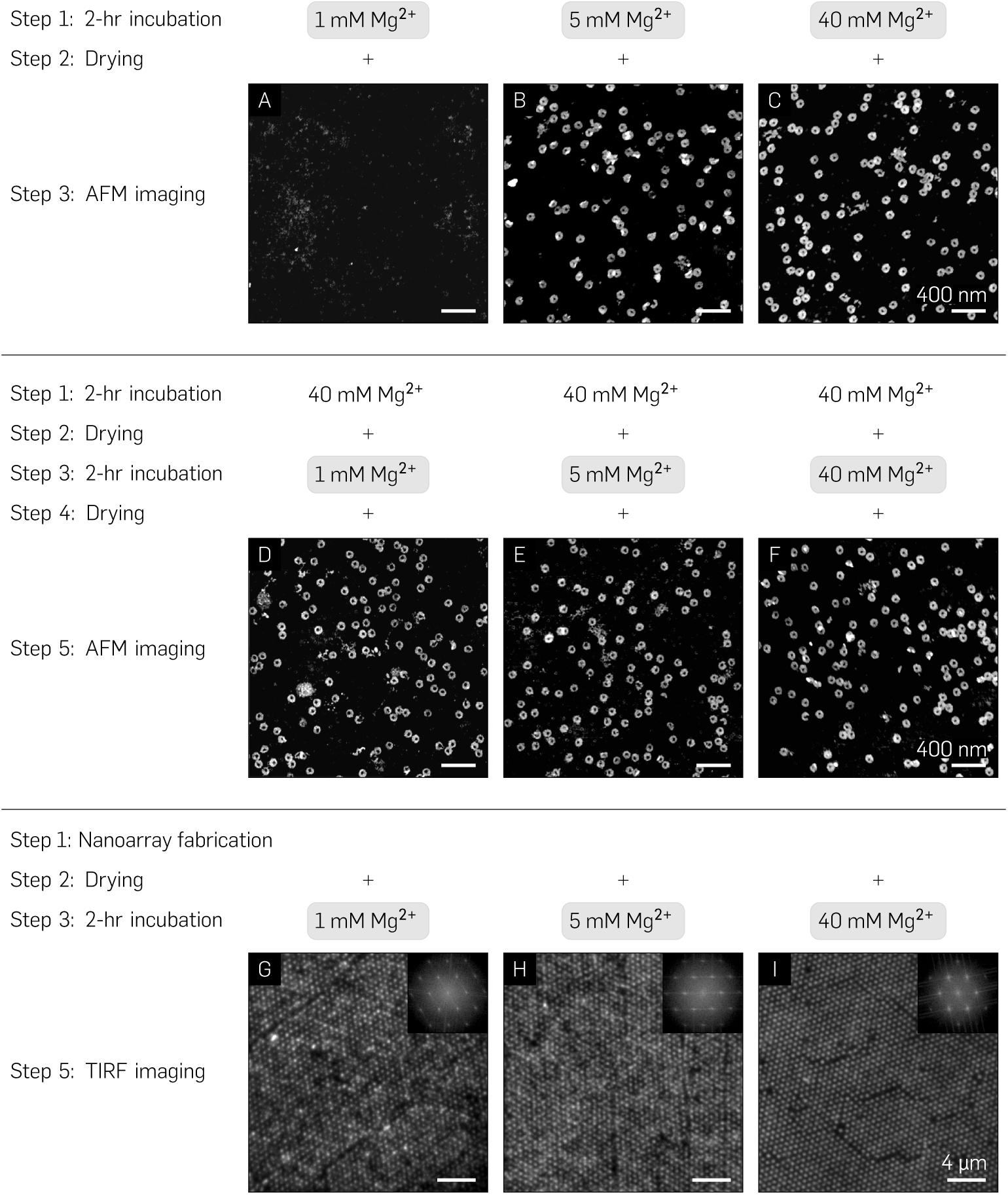
Nanoarray platform sensitivity to biologically-relevant multivalent cation concentrations. (**A**–**C**) AFM images of origami immobilization post-ethanol drying and rehydration on an activated glass substrate through direct incubation in 1, 5, and 40 mM Mg^2+^ buffer. (**D**–**F**) Incubation in 40 mM Mg^2+^ followed by ethanol drying and 2 hr rehydration in 1, 5, and 40 mM Mg^2+^, and consequent drying. (**G**–**I**) TIRF images and their corresponding FFTs (insets) of Mg^2+^–mediated immobilization on the nanoarray platform with 700-nm pitch at 40 mM Mg^2+^ followed by a 2-hour rehydration in 1, 5, and 40 mM Mg^2+^ buffers.

Previous DOP studies have demonstrated robustness at low salt conditions through multiple surface chemistry remodeling steps as well as covalent linkage of origami molecules onto already modified substrates-of-interest^36,37^. It is evident that surface chemistry forms the backbone of interactions between origami nanostructures and the substrate for DOP-based biophysical experiments. For simplicity and reproducibility, we optimized all experimental parameters for commercially available glass substrates routinely utilized for single-molecule biophysics experiments. With respect to the sensitivity of nanoarray to low divalent concentrations, we suspect that the ethanol drying process sequesters and stabilizes Mg^2+^ bridges between the origami and the silanol groups on the binding sites such that origami are conserved in an entropically-favorable energy state, mitigating dissociation or structural disintegration upon rehydration. Subsequent resuspension in lower divalent salt concentration has little-to-no adverse effects on these immobilized origami. To understand the physics underlying the drying process, additional experiments may be necessary. Most SMEs tend to last on the order of a few minutes-to-tens of minutes rather than a few hours, and would benefit from the robust nature of this platform. Further proof of functional and structural robustness is found in its long shelf life of several months post-drying at room temperature without the need for sophisticated storage (**Fig. S6**).

### Nanoarray platform facilitates optimal quality, high throughput, and deterministic single-molecule biophysics experiments

In order to quantify the efficiency of single-molecule incorporation prior to performing biophysics experiments on the nanoarray platform, we designed each origami baseplate to attach six fluorophore-labeled strands (in a hexagonal arrangement).We measured the incorporation efficiency of the designed strands to be 56%^60^ (**Fig. S7**, and section **S7**). Following the photobleaching experiment, we used the optical characterization technique of DNA-PAINT to benchmark the accessibility of biomolecules on the nanoarray platform (**Fig. 1C**). NSL-based placement positions a single origami nanostructure in a diffraction-limited area 74% of the time (**Fig. S8**) as opposed to traditional DNA PAINT experiments that rely on randomly-deposited DNA origami^61^. We used particle averaging, previously utilized in DNA-PAINT studies^9,11^, as a means to improve image resolution and compare control experiments (stochastic immobilization) against nanoarray-based experiments (deterministic immobilization). We first provide AFM images as evidence that even at low concentrations of origami (100 pM; **Fig. 5A**), it is likely that 2 or more structures could co-localize in a diffraction-limited spot. An increase in concentration to improve throughput results in a higher fraction of structures overlapping each other (500 pM; **Fig. 5B**). However, when patterned on a glass substrate by a distance slightly greater than the diffraction-limit (**Fig. 5H**), up to 74% of origami molecules singly occupy individual binding sites (**Fig. 3O**). We arranged three “docking” strands per vertex of a hexagon (**Fig. S9**) to counteract the low conjugation efficiency and transiently bind fluorescently-labeled “imager” strands in solution. Control experiments with randomly dispersed origami were first performed to justify conducting DNA-PAINT on origami immobilized through Mg^2+^-bridges on activated, and/or dehydrated glass coverslips. In addition to the HMDS layer (**Fig. 5D**), which is intrinsically part of the nanoarray fabrication process, we passivated the glass surface against non-specific interactions of fluorescent, *ss*DNA via a 0.05% (v/v) Tween-20 detergent^62^ in the 40 mM Mg^2+^, Tris-HCl “placement” buffer, pH 8.3 (**Fig. 5E**). In the absence of Tween-20 passivation, a honeycomb lattice corresponding to single-stranded imager strands interacting non-specifically with the background was observed (**Fig. 5F**). Therefore, by facilitating specific interactions with the probe strands on origami nanostructures in the binding sites, Tween-20 passivation aided in improving the Signal-to-Noise ratio (SNR). This technique of passivation enabled DNA-PAINT imaging quality on the nanoarray platform comparable to that routinely reported with PAINT studies using standard imaging and data processing protocols^11^ (**Fig. 5G** and **Movie A**). High-density PAINT experiments were subsequently performed on patterned substrates with inter-origami pitches of 350-nm to provide mostly single origami per binding site and maintain diffraction-limited resolvability of grids. A fluorescence micrograph of the patterned dataset (350-nm, 400 pM patterned) is also presented here (**Fig. 5G**).

**Fig. 5.**
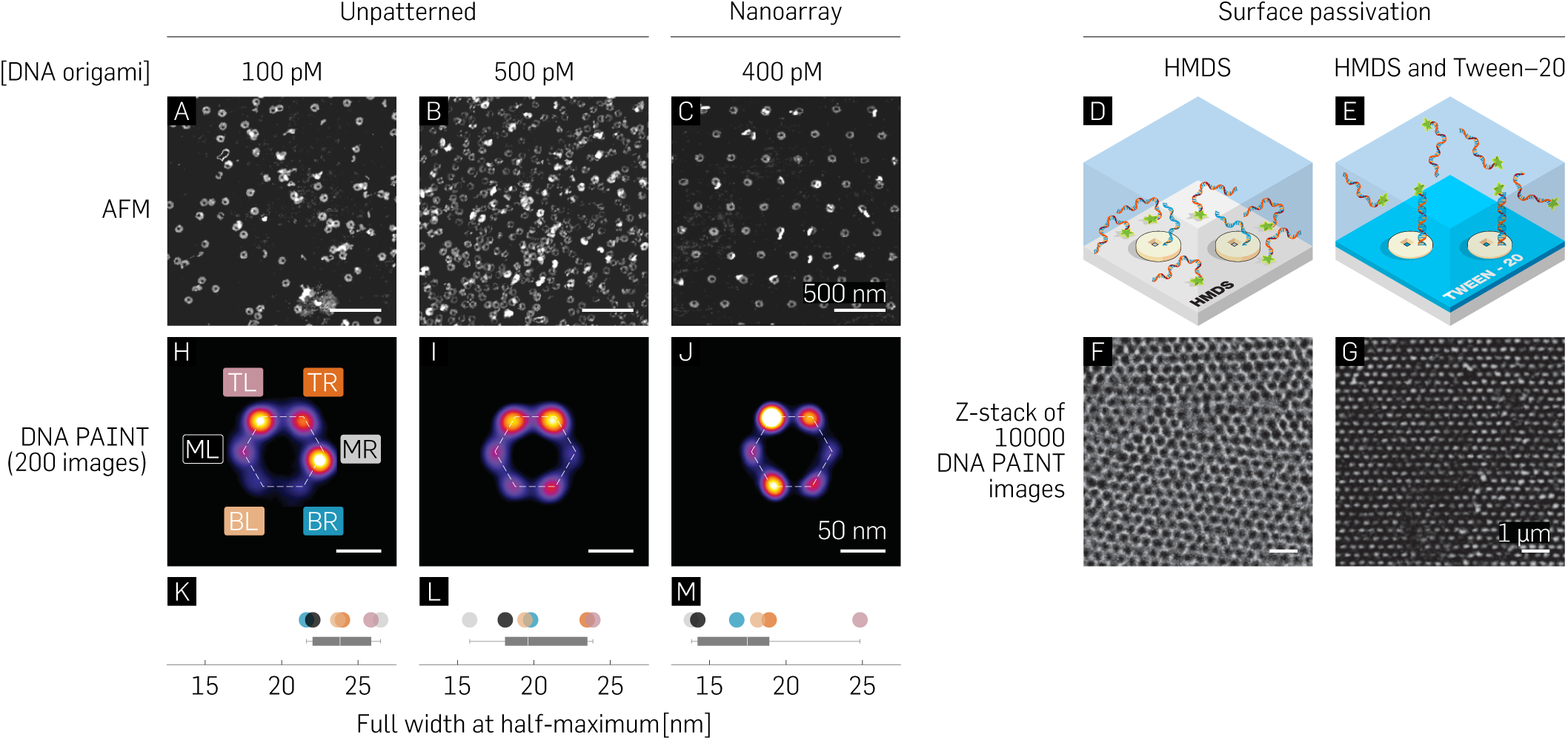
(**A**–**C**) AFM images contrasting stochastic single-molecule immobilization for low (100 pM), and high (500 pM) origami concentrations with origami deterministically patterned at the diffraction limit. (**D**–**G**) Schematic representation and experimental results from DNA-PAINT studies with (**D** and **F**) and without (**E** and **G**) Tween-20 treatment. (**H**–**M**) Averaged images of 200 manually-picked structures for low and high concentrations of stochastically immobilized origami (**H** and **I**), and patterned origami (**J**); and their respective full widths at half-maximum (**K**–**M**.)

Individual structures were averaged using the image processing software, Picasso^11^ (**Fig. 5H–J**), and their full width at half maximum (FWHM) measured as a metric to characterize the point spread function (PSFs) for the “sum” image. A standard analysis pipeline in Picasso comprises drift correction followed by manual or automatic single particle selection/picking, and finally, particle averaging^11^. We performed PSF comparisons (**Fig. 5K–M**) between the low (100 pM), and high concentrations (500 pM) of randomly immobilized origami with patterned origami (400 pM) for manually-picked structures (**Fig. 5H–M**) and automatically-picked structures (**Fig. S10**). We note that all origami used for these experiments broke up-down symmetry (20-T staple strands) and were therefore expected to have specific interactions with the imager strands. We also present a fluorescence micrograph of an exemplary patterned PAINT-dataset of 11,000 frames (350-nm) collapsed along the Z-axis prior to analysis (**Fig. S8**). To generate an averaged image, Picasso allows manual picking of structures (**Fig. 5H–J** or automatically picks structures similar to an initial user input of 5-10 structures (**Fig. S10**)). The averaging process of multiple structures involves aligning using their center of mass and finally through translational and rotational alignment over multiple iterations.

We picked 200 structures with ≥ 4 vertices present for the averaged image in each dataset (**Fig. 5H–J**). While the manually-picked structures from the randomly-immobilized origami had PSFs (Point Spread Function; mean ± SEM) of 23.94 ± 1.69 nm (low), and 20.07 ± 1.42 nm (high), the patterned data exhibited a slightly lower average value of 17.78 ± 1.26 nm (**Fig. 5K–M**). For automatically picking-based averaging, it is plausible that multiple overlapping structures in the high concentration case confound the software’s ability to accurately localize individual origami structures and their associated docking strands, ultimately leading to a degradation in averaged image quality (**Fig. S10**). We hypothesize that the bright points distinctly visible in both, automatically-as well as manually-picked structures could be a combination of two factors: location/sequence-dependent strand conjugation efficiency, as well as random noise. We expect random noise to be a factor especially in the case of automatically-picked structures with its source being non-specific interactions with– the surface, deformed origami, multiple overlapping origami, or gold nanoparticles used as fiducial markers for drift correction. For the manually-picked structures, we expect that the symmetry of the hexagonal pattern contributes to the software localizing certain vertices more brightly than others. This may be due to uncontrollable parameters (scoring function, alignment precision, etc.) in the analysis pipeline as well as the low occurrence of structures containing all six vertices.

As a quality control check to ascertain that the bright vertices were not solely a random function of the software analysis and might indicate a probability of certain strands being conjugated/accessible more than others, we excluded the “docking” strands for two vertices (**Fig. S9**). We observed that there was a low occurrence of at least one of the locations along the horizontal axis of the origami. Another possible reason for this could be the rotational symmetry along the vertical axis biasing the software rotational alignment to make one vertex brighter than the other. Regardless, our observations provide evidence that the nanoarray platform could serve as a potential solution to the concentration *vs*. throughput conundrum without compromising data quality. Furthermore, due to its intrinsically deterministic nature, it is amenable to software automation for simpler data analysis paradigms. These advantages of the platform could be leveraged for the benefit of a myriad of SMEs such as single-molecule FRET^7^.

## Conclusion

In summary, we have developed a cleanroom-free, DNA origami placement technique which surpasses the single-molecule binding efficiency imposed by Poisson statistics on traditional single-molecule deposition methods. The technique circumvents the need for sophisticated equipment and training previously required for fabricating single-molecule nanoarrays on the meso-to-macro-scale; all at ∼ $1 per chip. We characterized binding site sizes concomitant with various nanosphere diameters via AFM and EM. This provides a framework for the programmed placement of appropriately sized 2D or 3D DNA nanostructures for various single-molecule applications on addressable glass substrates. We report that a nanosphere diameter of ∼ 350 nm is essential to optimize the binding, and diffraction-limited imaging of single, circular DNA origami nanostructures (∼ 75%) and their associated payloads on high-density grids. We validate the robustness of this platform for *in vitro* single-molecule experiments under low divalent salt concentrations and demonstrate a shelf life of up to 10 months. We demonstrate the high-throughput and deterministic single-molecule experiments such as super-resolution, traditionally stochastic, DNA-PAINT without compromising data quality. We envision that the platform will be of great utility to biophysics, protein biochemistry, and digital diagnostics owing to its ability to democratize maximum throughput single-molecule experiments with bench-top fabrication in any conventional laboratory setting.

## Acknowledgements

We thank M. Kennedy and E. Le for support with data collection, and H. Sasaki, A. Auer, and R. Jungmann for helpful discussions on DNA–PAINT. This work was supported by National Institutes of Health Director’s New Innovator Award (1DP2AI144247) to RFH and Arizona Biomedical Research Consortium (ADHS17-00007401) to RFH; Office of Naval Research (N00014-17-1-2610 and N00014-18-1-2649) to PWKR and National Science Foundation (CCF-1317694 and CMMI-1636364) to PWKR. AFM data were collected in the lab of H. Yan at Arizona State University. SEM images were acquired at the Center for Solid State and Electronics Research at Arizona State University.

## Supporting Information

### S1. Materials and Methods

#### DNA origami design, preparation and purification

##### caDNAno file and supplementary files

The caDNAno design file, list of staples, a staple map, as well as a supplementary movie of raw DNA–PAINT data are included as a zip archive: `Origami designs+staples+movie.zip`.

### S2. Materials and Methods

#### Design

A circular origami with a square hole was designed using caDNAno http://cadnano.org/ as detailed by Gopinath *et al*.^59^. To control the face of the origami that binds to the binding site, we position all staple ends on the same face of the origami so that single-stranded 20T extensions to 5^*I*^ staple ends would all project from the same face of the origami.

#### Preparation

Staple strands (Integrated DNA Technologies, 640 nM each in water) and the scaffold strand (single-stranded p8064, 100 nM from Tilibit) were mixed together to target concentrations of 100 nM (each staple) and 20 nM, respectively (a 5:1 staple:scaffold ratio) in 40 mM Tris, 20 mM Acetate and 1 mM Ethylenediaminetetraacetic acid (EDTA) with a typical pH around 8.6, and 12.5 mM magnesium chloride (MgCl_2_) (1x TAE/Mg^2+^). 100 µL volumes of staple/scaffold mixture were heated to 90 °C for 5 min and annealed from 90 °C to 25 °C at 0.1 °C/min in a PCR machine. We used 0.2 mL TempAssure™tubes (USA Scientific). Once purified, the origami were stored in 0.5 mL DNA LoBind tubes (Eppendorf) to minimize loss of origami to the sides of the tube.

For annealing DNA–PAINT origami, and any other origami annealed overnight, a ramp of 0.004 °C/min was used in the critical “folding” range of 60–50 °C, a 0.005 °C/min ramp was used between 70–60 °C and 50–40 °C, and a 0.1 °C/min ramp was used between the 90–70 °C and 40–25 °C temperature ranges. “Docking strand” staples were introduced at 75–100x excess to the annealing mix.

#### Purification

A high concentration of excess staples will compete with DNA origami and inhibit DNA origami placement. Thus, origami were purified away from excess staples using 100 kD molecular weight cut-off filters (MWCO) spin filters (Amicon Ultra-0.5 Centrifugal Filter Units with Ultracel-100 membranes, Millipore, UFC510024). By the protocol below, recovery is generally 40–50% and staples are no longer visible by agarose gel electrophoresis:

1. Wet the membrane of the spin filter by adding 500 µL 1x TAE/Mg^2+^.
2. Centrifuge at 6000 rcf for 5 min at room temperature (RT), until the volume in the filter is ∼ 80 µL.
3. Discard the filtrate.
4. Add 100 µL of unpurified origami and 300 µL 1x TAE/Mg^2+^. Spin at 6000 rcf for 5 min at RT.
5. Discard the filtrate.
6. Add 420 µL 1x TAE/Mg^2+^ and spin at 6000 rcf for 5 min at RT.
7. Repeat step (4) two more times.
8. Invert the filter into a clean tube and spin at 6000 rcf for 5 min at RT to collect purified origami (∼80 µL).

Note In case of DNA origami annealed with a 75–100x excess of fluorophores (for photobleaching) or DNA–PAINT, spin the filter at 2000 rcf for 15 min, 5–7 times before inverting into a new tube to collect the purified product. This is to avoid fluorophores (and associated origami) from sticking to the sides of the filter and adversely affecting the purified origami yield or causing origami aggregation/deformation. Always check the purity of origami using agarose gel electrophoresis (100 V, 1%, 1x TAE, 1 hr).

The total time required for this purification is roughly 30–120 min. Post-purification, origami are quantified using a NanoDrop spectrophotometer (Thermo Scientific), estimating the molar extinction coefficient of the DNA origami as that of a fully double-stranded p8064 molecule (extinction coefficient = 164,568,055 /M/cm). We typically work with stock solutions of 20–30 nM DNA origami (3–5 OD). The working concentration for origami during placement is 100–500 pM, which is too small to be measured with the NanoDrop. Single-origami occupancy is sensitive to origami concentration, therefore, to maintain consistency for each series of experiments, a single high concentration stock solution (from a single purification) was made and diluted to 100–500 pM as needed. Origami concentration was optimized for best placement results for each origami stock.

### S3. Fabrication of binding sites and origami placement

#### Materials and equipment required

1. 10×10 mm^2^ coverslips (Ted Pella, 260375-15).
2. Plasma cleaner (Harrick Basic Plasma Cleaner PDC-32G/PDC-32G-2)
3. Hotplate and stirrer (Denville)
4. Desiccator (Hach, Product no. 223830)
5. Branson ultrasonic bath, AFM (Bruker FastScan).
6. Appropriately sized Polystyrene (PS) microspheres (3000 Series Nanosphere; Size Standards (4000 Series Monosized 1 µm particles 4009A; 700 nm [3700A]; 495 nm [3495A]; and 400 nm [3400A]), Thermo Fisher Scientific).
7. Passivation agent: HMDS (440191–100 mL, Sigma).

#### Protocol for binding site creation

1. Isopropanol (IPA) wash for 2 min.
2. Blow dry glass chip with nitrogen.
3. 10-min air plasma cleaning in Harrick plasma cleaner at ∼ 18 W (“High” setting).
4. In an eppendorf tube, pour 10 drops (∼ 360 µL suspension) of 1 µm/700 nm/500 nm/400 nm PS nanospheres. Gently vortex the nanospheres before use.
5. Spin at 8,000–10,000 rpm for 5 min faster and/or longer spinning for smaller nanosphere sizes).
6. Remove supernatant and add 360 µL of ultrapure water to re-suspend pellet.
7. Spin at 8,000-10,000 rpm for 5 min.
8. Remove supernatant and resuspend pellet in 25% ethanol and 75% water (∼ 3.5x more concentrated, *i.e.* 100 µL). Pipette/vortex aggressively to resuspend all particles (∼ 6.5*e*^10^ particles/mL for 1 µm nanospheres at 1% w/w solids).
9. Drop-cast onto activated chip surface and let dry at ∼ 45° angle at R.T (resting against a glass stirrer or similar object). Cover entire surface (generally requires 5-10 µL for a 10 *×* 10 mm^2^ chip). Once dried, you should be able to observe a diffraction pattern (crystalline structure) confirming the existence of a close-packed monolayer/multilayer of nanospheres. If unsure, check under a microscope.
10. Heat at 60 °C for 5 min to remove any moisture.
11. 2-min “descum” plasma in air at ∼ 18 W in Harrickplasma cleaner.
12. In a desiccator, add 8–10 drops of HMDS (in a glass cuvette), and deposit under a vacuum seal for 20 min. This should work equally-well in an enclosed petri dish.
13. Lift-off PS nanospheres with water sonication in a Branson ultrasonic bath for 30–60 sec to create origami binding sites. In the absence of an ultrasonic bath, continuous stirring in water for a longer period of time is adequate. The nanospheres visibly come off the surface.
14. Blow dry with a nitrogen “gun”.
15. Bake at 120 °C for 5 min to stabilize the HMDS on the surface.

Note If you find areas without patterned DNA origami or binding sites, you may need a higher concentration of nanospheres (this is generally observed for 700–1000 nm nanospheres). A good sanity check is to label the origami with fluorophores, if possible, and observe under a fluorescence microscope for grids. AFM can sample a small fraction of a chip surface. For <500 nm nanosphere sizes, finding origami grids should not be a problem.

#### Origami placement experiments

1. Thermal Cycler (Life Technologies) for origami annealing.
2. 100 kDa spin filter columns (Amicon).
3. A benchtop centrifuge (Denville, 6000 g, 3–5 rounds of 5-min spin) for origami purification.
4. Origami: Modified circle with a square hole *aka* Death Star^59^.
5. Tris-HCl buffer (Buffer 1: pH 8.35, 40 mM Mg^2+^, 40 mM Tris, and Buffer 2: pH 8.9, 35 mM Mg^2+^, 10 mM Tris) [Magnesium Chloride Hexahydrate | M9272-500G, Sigma; Tris, T-400-1 GoldBio].
6. 50%, 75%, and 85% Ethanol (459836 Sigma Aldrich) in ultrapure water.

#### A step-by-step protocol for origami placement and washing steps

1. Incubate chips with ∼100–200 pM origami (nominal concentration for 1 µm pitch, concentration inversely proportional to nanosphere size) in ∼ 40 mM Mg, Tris-HCl (40 mM Tris) buffer (pH-8.3) for 60 min.
2. Wash in ∼ 40 mM Mg, Tris-HCl (40 mM Tris) buffer (pH-8.3) for 5 min either manually or automatically using a peristaltic pump or shaker in a petri dish.
3. Transfer to ∼40 mM Mg, Tris-HCl (40 mM Tris) buffer (pH-8.3) + 0.07% Tween 20 and wash for 5 min.
4. Transfer to ∼ 35 mM Mg, 10 mM Tris (pH-8.9) to hydrolyze HMDS and lift off origami non-specifically bound to the background and wash for 5 min.
5. For AFM characterization, transfer to ethanol drying series: 10 seconds in 50% ethanol, 20 seconds in 75% ethanol, 2 min in 85% ethanol.
6. Air-dry, followed by AFM/fluorescence verification of patterning.

Note All of the work reported in this paper was performed with spin-column purified origami, which is suitable for small amounts of origami. After purification and quantification, it is critical to use DNA LoBind tubes (Eppendorf) for storage and dilution of low concentration DNA origami solutions. Low dilutions, e.g. 100 pM, must be made fresh from more concentrated solutions and used immediately— even overnight storage can result in total loss of origami to the sides of the tube. Addition of significant amounts of carrier DNA to prevent origami loss may prevent origami placement, just as excess staples do. We have not yet determined whether other blocking agents such as BSA might prevent both origami loss and preserve placement.

### S4. AFM characterization

All AFM images were acquired using a Dimension FastScan Bio (Bruker) using the “short and fat”, or “long and thin” ScanAsyst-IN AIR or ScanAsyst-FLUID+ cantilever (“sharp nitride lever”(SNL), 2 nm tip radius, Bruker) in ScanAsyst Air or Fluid mode. All samples were ethanol dried prior to imaging. Single and multiple binding events for placed origami were hand-annotated for origami occupancy statistics and image averaging of arrays (imageJ) was used to determine binding site size.

### S5. SEM characterization

Images of close-packed nanosphere crystals, as well as individual nanosphere cross-sections were obtained using a Hitachi S-4700 Field Emission Scanning Electron Microscope (ASU Nanofab, Center for Solid State Electronics Research, Tempe, AZ) at 1–5 keV and the stage (or electron beam) was manipulated as required. In order to prevent charging effects and distortion of the image collected, a sputter coater (Denton Vacuum Desk II, New Jersey) was used to coat the specimen (glass with nanospheres) with Gold-Palladium (Au-Pd), and carbon tape was used to provide a conduction path from the glass surface to the SEM stub (ground). For the cross-sectional images specifically, the glass coverslip was broken in half post sputter-coating and wedged inside a standard cross-sectional SEM sample holder such that the electron beam impinged directly on the flat edge of the glass coverslip to visualize the contact areas between the nanospheres and the glass surface. Measurements from high-resolution images were made manually using imageJ.

### AFM *vs*. SEM analysis of binding site size

Over the range of nanosphere diameters tested, we found a global discrepancy of ∼ 11% between the linear fits, with SEM (*a* = 0.38*d*_*ns*_; **Fig. 3A–F**) providing consistently larger estimates than AFM (*a* = 0.27*d*_*ns*_ ; **Fig. 3G–L and N**). We first note that each of the SEM mean and SD values are gleaned from *N* =≤ 10 nanospheres, whereas corresponding AFM values are determined using weighted means and SDs from averaged images of *N >*400 binding sites for any given nanosphere diameter.

We offer two possible explanations for the observed discrepancy:

i. HMDS is a miniscule molecule which can lead to larger coverage of interstitial spaces between nanospheres than can be accurately measured using an indirect technique such as SEM. This may result in overestimation of masking areas when examining electron micrographs. In addition, all SEM images were collected by sputter coating sample cross-sections with a ∼ 10-nm Gold-Palladium (AuPd) layer (for conductivity) which may further contribute to higher estimated values. AFM, however, provides a direct mode of measurement post-passivation with HMDS, and can facilitate a more precise estimation of the “footprint” of each individual nanosphere in an HCP layer.
ii. On closer observation of electron micrographs, we found an apparent distortion of nanosphere geometry along the *xy*-axes in keeping with the phenomenon of Hertzian contact (**Fig. 3A**, top row). This alteration in morphology could be linked to the position of each nanosphere, and/or the electron beam with respect to the substrate edge, as well as the relative position of each nanosphere to other, adjacent spheres, and the physical contact between individual nanospheres and the glass surface. It is plausible that attractive forces during the solvent evaporation process contribute to the departure from a spherical to a more flattened shape upon interaction with neighboring colloidal particles. Using a simplistic deformation model to support this hypothesis **Fig. S2**, we note that for an 8% distortion along the vertical axis of a nanosphere (80 nm for a 1 µm diameter), the predicted binding site size follows the experimental SEM values closely. This may help explain the discrepancy between the observed SEM and AFM values in a quantitative manner, with SEM images overestimating the effect of distortion caused by physical deformation on binding site sizes obtained.

### S6. DNA–PAINT

All TIRF experiments were conducted on a benchtop super-resolution Oxford Nanoimager (Oxford, UK). For control DNA–PAINT experiments, a glass chip was activated for 10 min, followed by the creation of a “flow chamber” (using double sticky Kapton polyimide adhesive tape; Amazon) and 30-min incubation of 400 pM DNA origami at 40 mM Mg^2+^. Non-specifically bound origami were washed off using several rounds of wicking the incubation buffer through the chamber. Next, a 0.05% Tween–20 (Cat no. P1379, Sigma Aldrich) v/v in 40 mM Mg^2+^ placement buffer was flown through several times before incubating the solution for 5 min. This prevents non-specific ssDNA binding during the experiment (**Fig. 5E and G**). Subsequent washing in Tween–buffer, and placement buffer was followed by the introduction of up to 5 nM P1-imager solution in placement-Tween buffer, 10x dilution of 40-nm Gold nanoparticles (fiducials for drift correction, Sigma Aldrich 741981), and an oxygen scavenging system (2x, 3x, 5x concentrations of PCA, PCD, and Trolox-Quinone respectively). To ensure the gold nanoparticles settle on the bottom chip, it was taped to a 96-well plate holder in a centrifuge and spun at 150 g for 5 min, ensuring that the inlet and outlet of the flow chamber were sealed prior to spinning. Experiments with patterned chips were conducted by sticking the 10 *×* 10 mm^2^ coverslip onto a double-sided sticky Kapton tape and repeating the procedure as outlined above starting with incubation with 0.05% Tween–20 in placement buffer.

DNA–PAINT data were analyzed using Picasso^11^. Briefly, a dataset <4 GB was prepared for analysis on Picasso `Localize`. A minimum net gradient of 5,000 was chosen to avoid non-specific signals from being analyzed. After the fits were found, the `.hdf5` file was loaded into the Filter module of Picasso. This module allows localization precision filtering, as well as the filtering of “double” localizations by manually selecting a Gaussian profile of localization photons. The filtered localizations dataset is then loaded into Picaso `Render` where multiple cross-correlation-based drift corrections and multiple corrections based on picking fiducial markers on the sample 40-nm gold beads and/or origami themselves [“`pick similar`”]) were used to perform more precise drift correction. The threshold was adjusted prior to automatic or manual picking of structures to be averaged. The picked localizations were then registered into Picasso `Average` where they were aligned using center of mass followed by multiple iterations of rotational and refined translational alignment to form the final “summed” image. A nominal oversampling value of 200 was used to represent the structures prior to measuring the PSFs in imageJ using an ROI drawn around each vertex and finding its full width at half maximum (FWHM **Fig. 5K–M**).

**Table S1.** Buffer composition of DNA–PAINT experiments.

| Buffer components | Volume (μL) |
| --- | --- |
| Imager “P1” strand in 40 mM Mg<sup>2+</sup> + 0.05% Tween-20 (10 nM, stock) | 30 |
| 40 mM Mg<sup>2+</sup> + 0.05% Tween-20 | 16.7 |
| 40 nm gold nanoparticles | 6 |
| 50× PCA | 2.5 |
| 100× PCD | 1.8 |
| 100× Trolox–Quinone | 3 |
| **Total** | **60** |

Note

1. A concentration beyond 0.1% Tween–20 may result in disassociation of origami from the binding sites.
2. All datasets analyzed had duty cycles of 1:10–1:50 (10,000–12,000 frames) and were screened based on amount of photobleaching as a quality-control measure.

### S7. Photobleaching experiments

For the photobleaching experiments, we analyzed the intensity traces of origami molecules in response to laser excitation and observed steps corresponding to independent, stochastic fluorophore quenching events. Based on a histogram of the number of fluorophores experimentally found to be incorporated per origami baseplate we calculated the strand conjugation efficiency to be 56%. For these experiments we assumed that the fluorophore-of-interest was indeed conjugated to the strand complementary to the six handle strands, and that it was not photobleached prior to experimental observation. Yields of 84% have been previously reported^13^, and we hypothesized our observation of lower yield could possibly be due to poor strand accessibility. It is important to note that the circular origami have been experimentally shown to break up-down symmetry using staples modified with 20T extensions that act as entropic brushes, with 95.6% origami facing the designed-side up^59^. While this is experimentally advantageous in terms of number of structures facing right side up, and consequently the amount of useful data collected, it is plausible that these 20T extensions may result in steric hindrances and poor accessibility of the strands-of-interest. We tested several conditions such as circumventing the dehydration step (to rule out the accessibility problem), increasing strand-excess, annealing time, and additionally, directly annealing the fluorophore-labeled complementary strands with the handle strands. We did not find any significant changes in incorporation efficiency with yields of ∼ 60% in all cases. Based on these observations, we posit that the sub-par conjugation efficiency may be sequence-, strand concentration-, strand purity-, strand position on origami-, or origami purification strategy-dependent. This low yield, while concerning, is a broader concern for the field of structural DNA nanotechnology itself and a comprehensive examination may require a more concerted effort by the community.

Photobleaching experiments were performed immediately after grid formation in imaging buffer (1x TAE. 12.5 mM Mg^2+^) and oxygen scavenging similar to DNA-PAINT). Control experiments of randomly immobilized origami were also performed. There was no difference in the conjugation efficiency with and without patterning. Similarly, no difference was observed when origami were pre-labeled (*i.e.* P1P4-Cy3B strand annealed with origami in a one-pot reaction) in comparison to being post-labeled once on the surface of the chip. In both cases, fluorophore-labeled strands were added at a concentration between 10–100 nM (*i.e.* 10–100x excess). Laser intensity was adjusted in order to have a slow gradient of fluorophore intensity bleaching to make step-counting easier. Steps were quantified using two methods: imageJ^63^, and iSMS^64^ (http://inano.au.dk/about/research-groups/single-molecule-biophotonics-group-victoria-birkedal/software/). For the latter, the field-of-view was cropped, and the two ROIs were aligned to count the number of steps distinctly.

The following were the criteria when analyzing photobleaching data:

1. Must bleach completely.
2. Must have signal of a single molecule, and not aggregates.
3. Must show ≤ 6 photobleaching steps.
4. Must have a consistent step size (quanta).

### S8. Guide to troubleshooting binding site creation and origami placement

| Problem | Likely cause(s) | Potential solution(s) |
| --- | --- | --- |
| Nanosphere pellet is absent | 1. Spin speed is too low to sediment beads out of solution. | 1. Spin speed/time required for pellet formation is inversely proportional to nanosphere size and should be adjusted accordingly.<br>2. Use a different rotor. |
| Nanospheres do not adhere to the glass surface. | 1. Improper plasma activation.<br>2. Improper ethanol concentration. | 1. Contact angle measurements should delineate the difference between “inactive” and “active” glass surfaces.<br>2. Start at a nominal, 25% ethanol, but 50% should give desired results.<br>3. Low nanosphere concentration. |
| No nanosphere close-packing observed. | 1. Traces of surfactant in the original solution prevents self-assembly of nanospheres.<br>2. Final ethanol suspension might have surfactant due to incomplete wash steps. | 1. Wash $\geq 2$ times and resuspend in pure water before preparing a final suspension in 25% ethanol at the appropriate nanosphere concentration.<br>2. Carefully aspirate supernatant from the end opposite to the pellet on the Eppendorf tube. |
| Nanospheres do not come off after sonication. | 1. Sonication intensity may be low.<br>2. Dehydration bake temperature may be too high. | 1. Use an 80×40 mm<sup>2</sup> Pyrex petri dish (3140; Cole-Parmer) for sonication and leave the dish in the bath for *geq* 5 mins if necessary<br>2. The glass transition temperature ($T_g$) of polystyrene nanospheres is ~80 °C and should not be exceeded to prevent nanospheres from losing their integrity. |
| *Alternative: Chips can be inserted in a petri dish containing an organic solvent such as Tetrahydrofuran (THF, 401757, Sigma) heated to 60 °C for 5–15 mins to dissolve beads. Care must be taken when handling THF, and to reduce evaporation owing to its volatility.* |  |  |
| Binding site variability is observed over the surface of the chip. | 1. Insufficient dehydration baking time results in “water puddles” around the nanosphere “footprints” preventing HMDS from depositing all around the circumference.<br>2. Nanospheres are not NIST-standards approved, *i.e.* large standard deviation in nanosphere size. | 1. Increase initial dehydration bake time to 10 min.<br>2. (i) Check with the manufacturer to confirm SD from the nominal size of nanospheres.<br>(ii) Measure the distribution of nanosphere size. |
| Binding sites are hard to find using AFM. | 1. AFM tapping force under threshold value.<br>2. Scanned area is devoid of nanosphere close-packing. | 1. Tune scanning parameters to rationally to image binding sites<br>2. Proceed to origami placement to simplify AFM imaging. Origami nanostructures provide better contrast.<br>3. Sample at least one area from each quadrant of the chip to determine the source of the issue conclusively. |
| Impurities on the surface | 1. Improper cleaning<br>2. Dirty plasma chamber.<br>3. Failure to perform “descum” plasma step, which can leave the particulate matter in the interstitial spaces. | 1. More aggressive cleaning required (*e.g.* RCA).<br>2. Clean plasma cleaner chamber with isopropanol, let dry, run it without a sample for an hour, clean again (checking for contaminant residues after every step).<br>3. Filter all buffers used during wash steps.<br>4. Ensure petri dishes are clean.<br>5. If using automated washing, always sterilize peristaltic pump tubing by running boiling hot water through it first, followed by a gradient of 100%, 75%, and 50% ethanol in pure water, and finally placement buffer just before use. |
| Origami bound non-specifically all over the surface. | 1. Poor HMDS quality.<br>2. Bake step for HMDS stabilization was accidentally skipped. | 1. Dehydrate the wafer by baking before and after HMDS monolayer formation.<br>2. Keep pH<9, preferably at 8.3–8.5 |
|  | 3. Improper washing and failure to remove weakly bound origami from HMDS background<br>- new buffers may be required | 3. Reduce $\text{Mg}^{2+}$ concentration, and compensate by increasing origami concentration/incubation time<br>4. Keep incubation between 30–90 minutes<br>5. Remove weakly bound origami with more aggressive Tween-20 washes (0.08%, 10-20 minutes).<br>6. Remove any hydrophobic moieties (*e.g.* fluorophore) from the origami. Note that HMDS is hydrophobic. |
| Low DNA origami occupancy. | 1. Old chip with inactive sites.<br>2. Low origami concentration.<br>3. Short incubation time<br>4. Low pH, or Mg concentration. | 1. Sites remain optimally active for $\lesssim 4$ hours post-plasma cleaning.<br>2. Always optimize origami concentration (*e.g.* 4 chips, 200–400 pM for 400-nm spacing). Use higher origami concentration, prepare fresh dilutions, and use immediately<br>3. Check buffer pH (8.3–8.5), prepare fresh if necessary, with 35 mM $\text{Mg}^{2+}$.<br>4. Add a verification step to ascertain appropriate binding site diameter post nanosphere lift-off using AFM. |
| High multiple DNA origami occupancy. | 1. High origami concentration.<br>2. Long incubation time.<br>3. Oversized feature | 1. Optimize origami concentration every time fresh stock is prepared.<br>2. Use an incubation time between 30–90 minutes<br>3. Add a verification step to ascertain appropriate binding site diameter post nanosphere lift-off using AFM<br>4. Check buffer conditions |
| High background binding: Origami (balled-up [white “blobs”] or otherwise) | If HMDS passivation is good, origami cannot bind flush against the surface, and improper washing can lead origami to come off partially. Subsequent ethanol washing leads to any origami not bound completely to ball up and appear as white blobs during AFM imaging. | 1. Dehydrate the chip before and after HMDS monolayer formation taking care to keep first dehydration step below $T_g$ of polystyrene nanospheres<br>2. Keep placement buffer pH between 8.3–8.5<br>3. Remove weakly bound origami with longer Tween-20 wash and/or increasing Tween-20 concentration (<0.1% to prevent origami in binding sites from falling off). |
| Origami fall off during ethanol drying. | Too much time spent in dilute ethanol (<80%). | Move quickly from low to high % ethanol. |
| Origami ball up into site during ethanol drying and corners are double height. | Origami project partially onto non-sticky HMDS surface. | Hydrolyze HMDS surface before drying by washing in pH 8.9–9.0 buffer to mitigate background binding. |
| Binding sites are filled up, but no defined origami shapes observed | Excess of staple strands. | 1. Check scaffold, and origami integrity using gel electrophoresis and AFM pre- and post-purification<br>2. Scaffold quality might be a problem |

### S9. Supplementary Figures

**Fig. S1.**
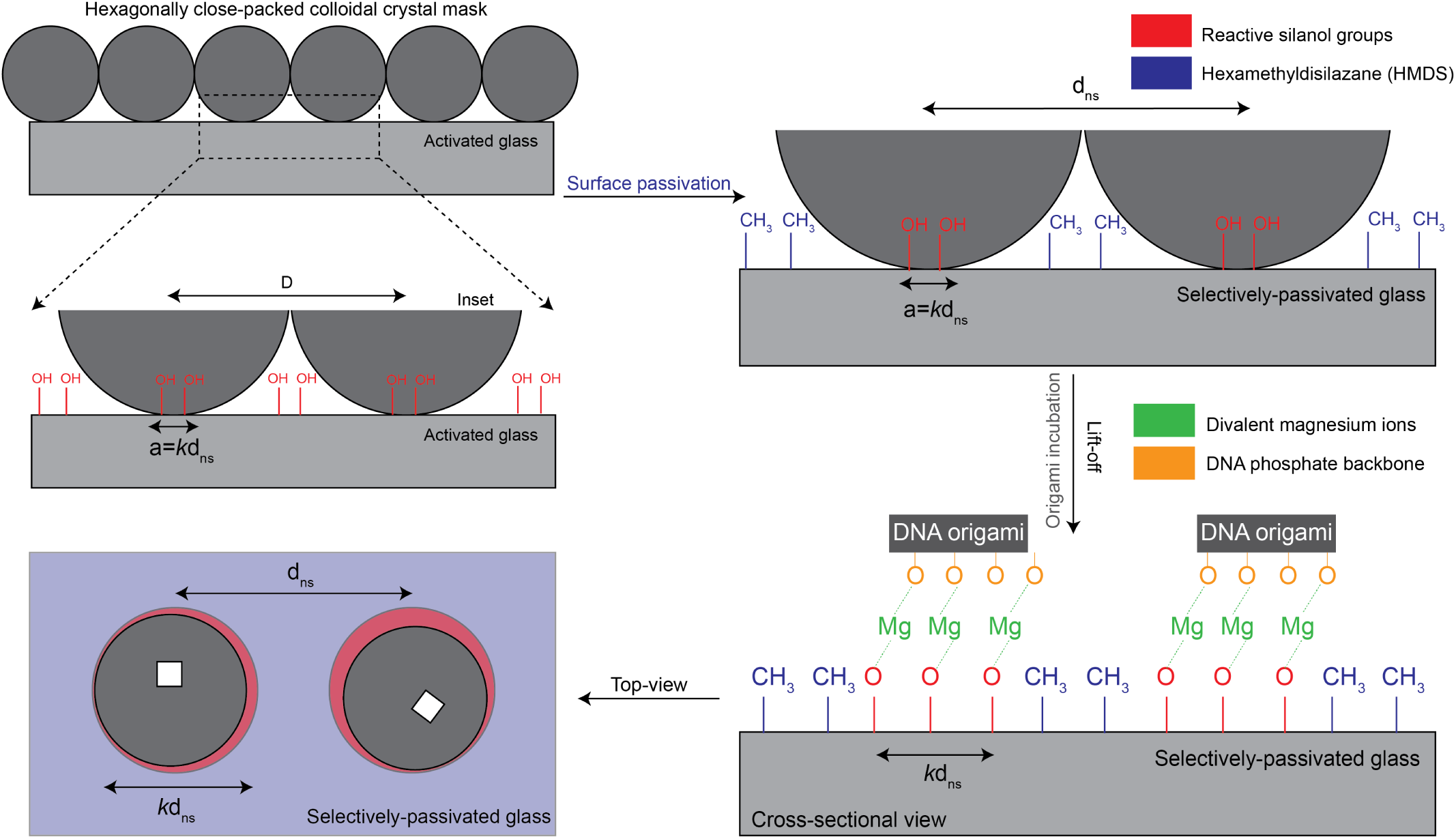
Schema of reactive silanol groups being protected by a colloidal crystal mask (CCM) of appropriately-sized nanospheres. Each nanosphere footprint is intrinsically linked to its diameter *a* by the relation: *a* = *kd*_*ns*_. A bulk surface passivation step with HMDS results in the selective passivation of the entire chip surface with neutral and hydrophobic methyl groups. Upon lift-off of nanospheres, magnesium-mediated placement of negatively charged DNA origami to the reactive silanol groups formerly protected by the spheres proceeds through a process of diffusion, and alignment, prior to immobilization^37^. The important parameters that determine quality of nanoarray formation are Mg^2+^ concentration, pH, incubation time, origami concentration, and adequate washing (to reduce spurious background bindings).

**Fig. S2.**
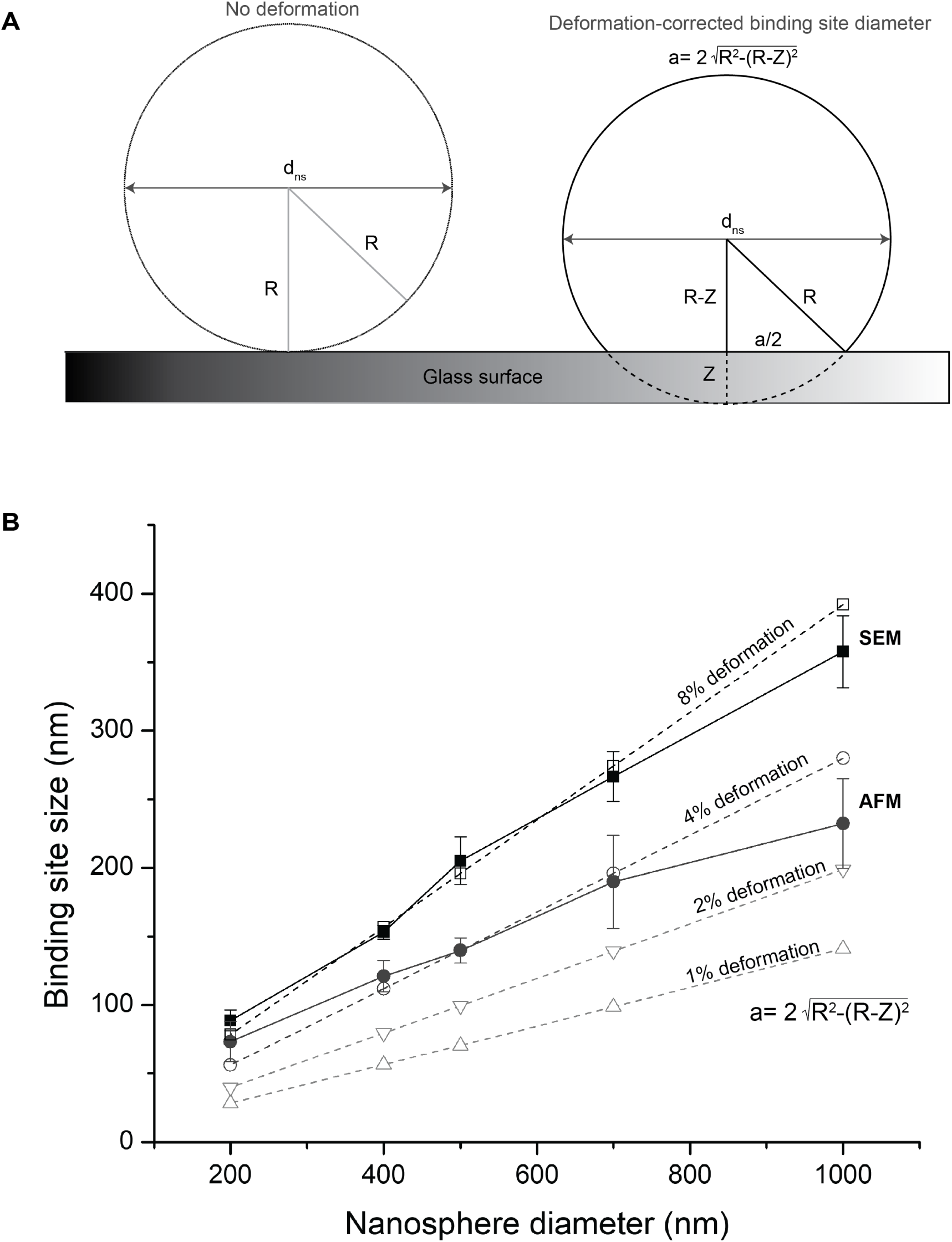
EM images (**Fig. 3A–F**) point to deformation of nanospheres resting on the surface due to Hertzian contact: interactions between adjacent spheres as a result of the capillary forces that drive them together during the process of close-packing, and interactions between the spheres and the surface. (**A**) The deformation-free case (left), and the distortion of nanosphere geometry owing to deformation (right), where *a* denotes the predicted binding site diameter owing to the deformation. (**B**) plots the deformation-associated binding site diameters (for 1%, 2%, 4%, and 8% deformation) in comparison with binding sites measured via SEM (**Fig. 3A–F**) and AFM (**Fig. 3G–L**). Based on the relationship provided here, the co-efficient *k* in the relationship *a* = *kd*_*ns*_ is 0.27 for AFM, and 0.38 for SEM, *i.e.* the binding sites are expected to be 27% and 38% of their corresponding nanosphere diameters. As seen here, a deformation of 8% closely follows the values obtained using SEM measurements (indirect) which overestimate the binding site size compared to the 4% deformation predicted by AFM (direct) values as alluded to in Supplementary Section **S5**.

**Fig. S3.**
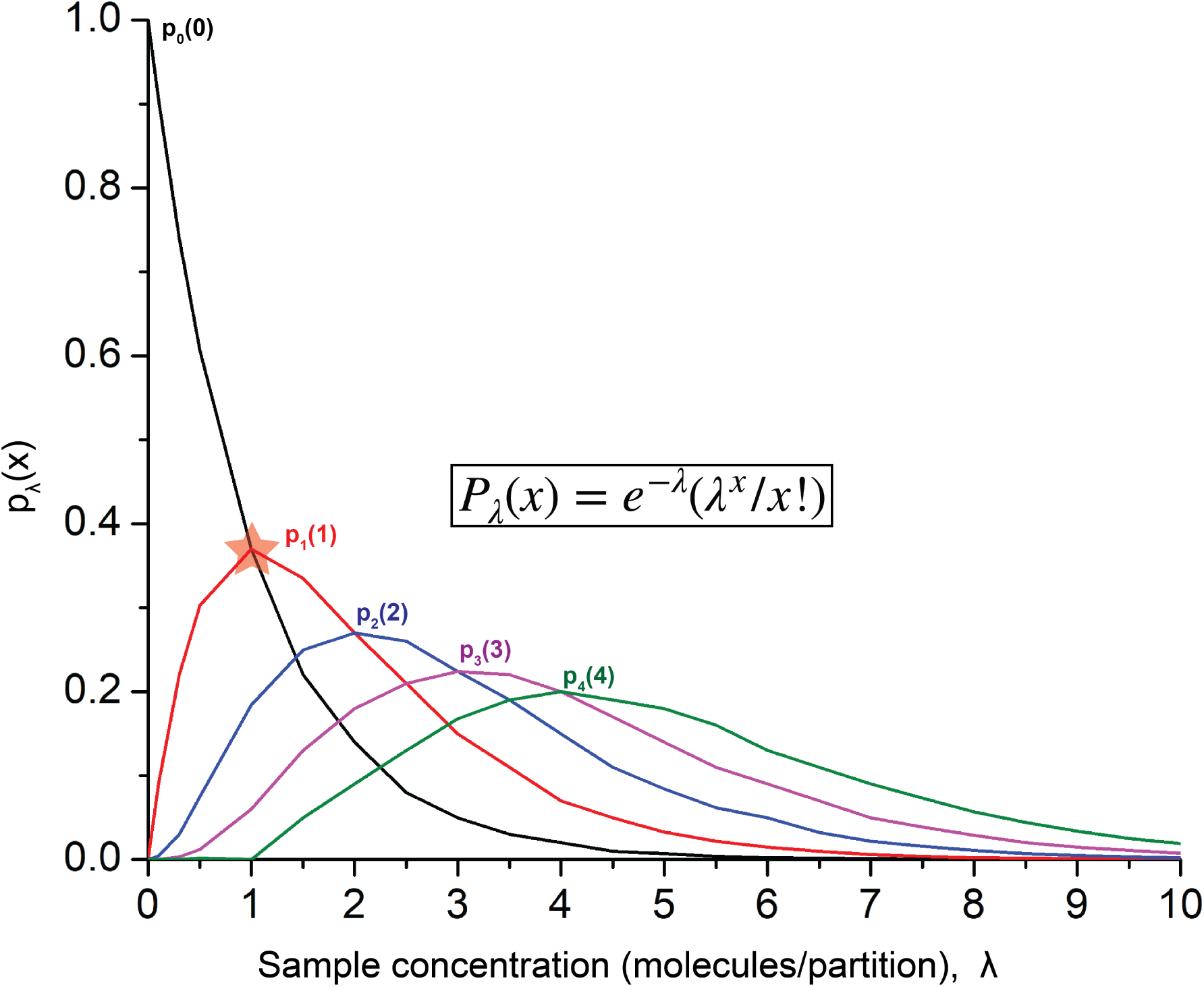
The Poisson distribution which poses a statistical limitation on the probability of a single molecule occupying in each partition/well on a substrate. It quantifies the probability for finding 0, 1, or more than 1 molecule in a single partition (binding site in this study) given a certain sample concentration. The highest single molecule occupancy/binding efficiency occurs in the case where the ratio of molecules to wells is one and is maximally 37% (red star). This is the case for every stochastic top-down loading process unless a steric hindrance approach in the form of a DNA origami macromolecule (in this study) is used to prevent multiple molecules from binding to the same spot, consequently driving the single occupancy beyond the Poisson limit as reported in (**Fig. 3O**; horizontal dashed line)

**Fig. S4.**
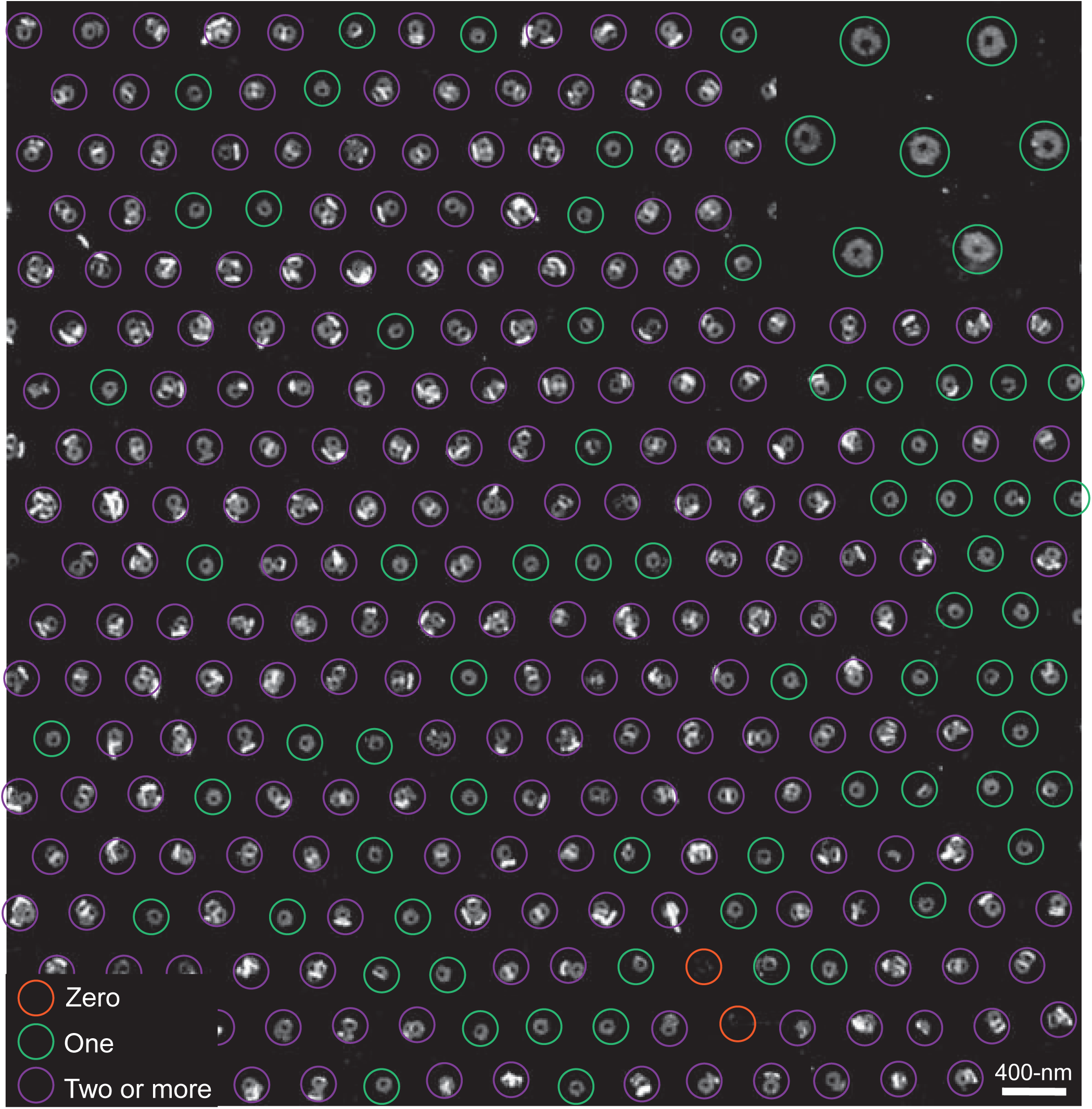
A representative AFM image of a DNA origami nanoarray fabricated using 15 mM Mg^2+^ by varying other governing global parameters such as origami concentration, incubation time, and buffer pH. Experimental parameters here were 250 pM origami, 90-min incubation, and pH = 7.8.

**Fig. S5.**
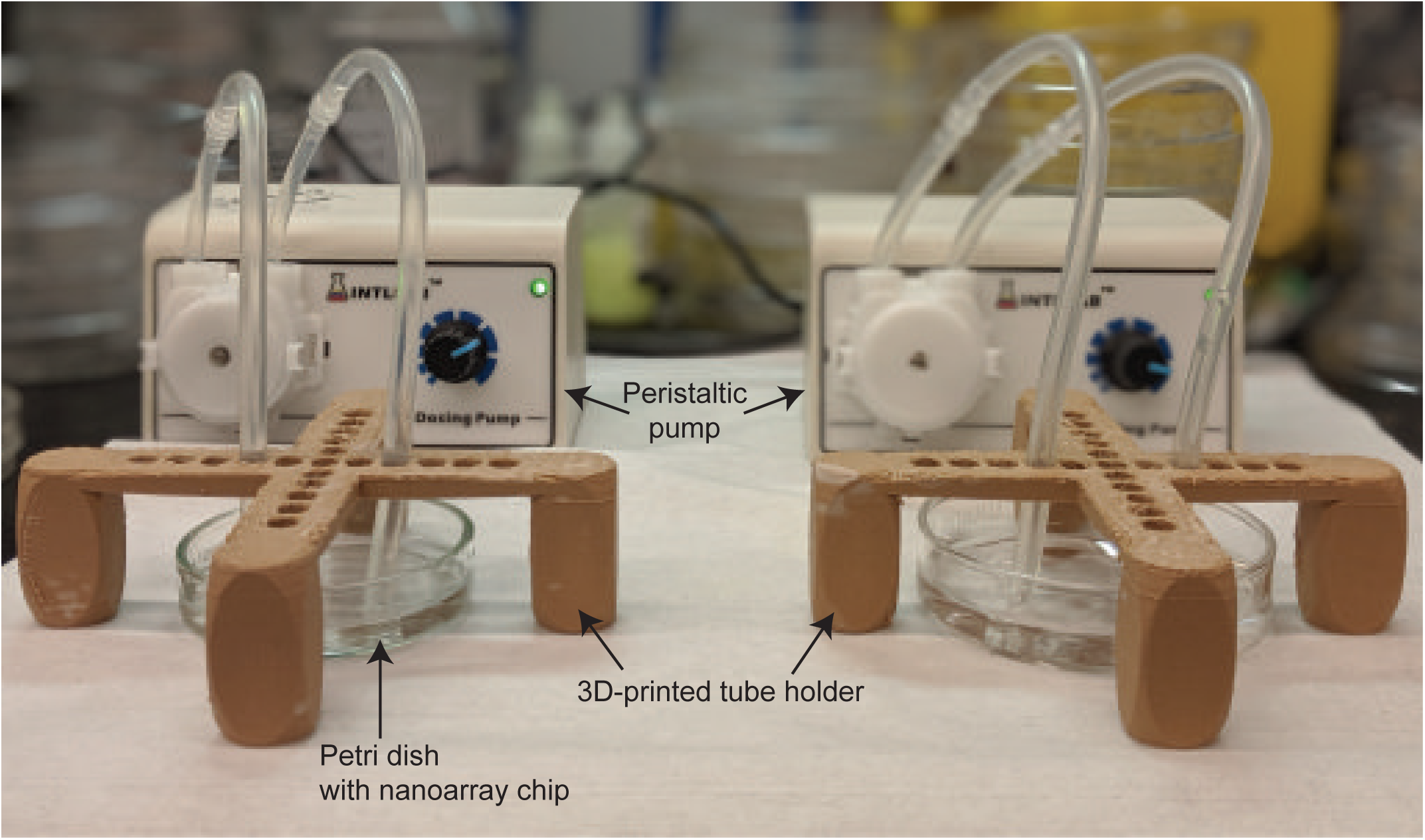
Peristaltic pumps for automating the three, 5-min wash steps for optimized cleaning of nanoarray chips. We use 3D-printed tubing holders to maintain a constant position for consistency in quality. The peristaltic pumps primarily mitigate user-variability introduced during the wash steps. Alternatively, a rocker with parameters adjusted for optimal washing can be used for consistent automated wash steps.

**Fig. S6.**
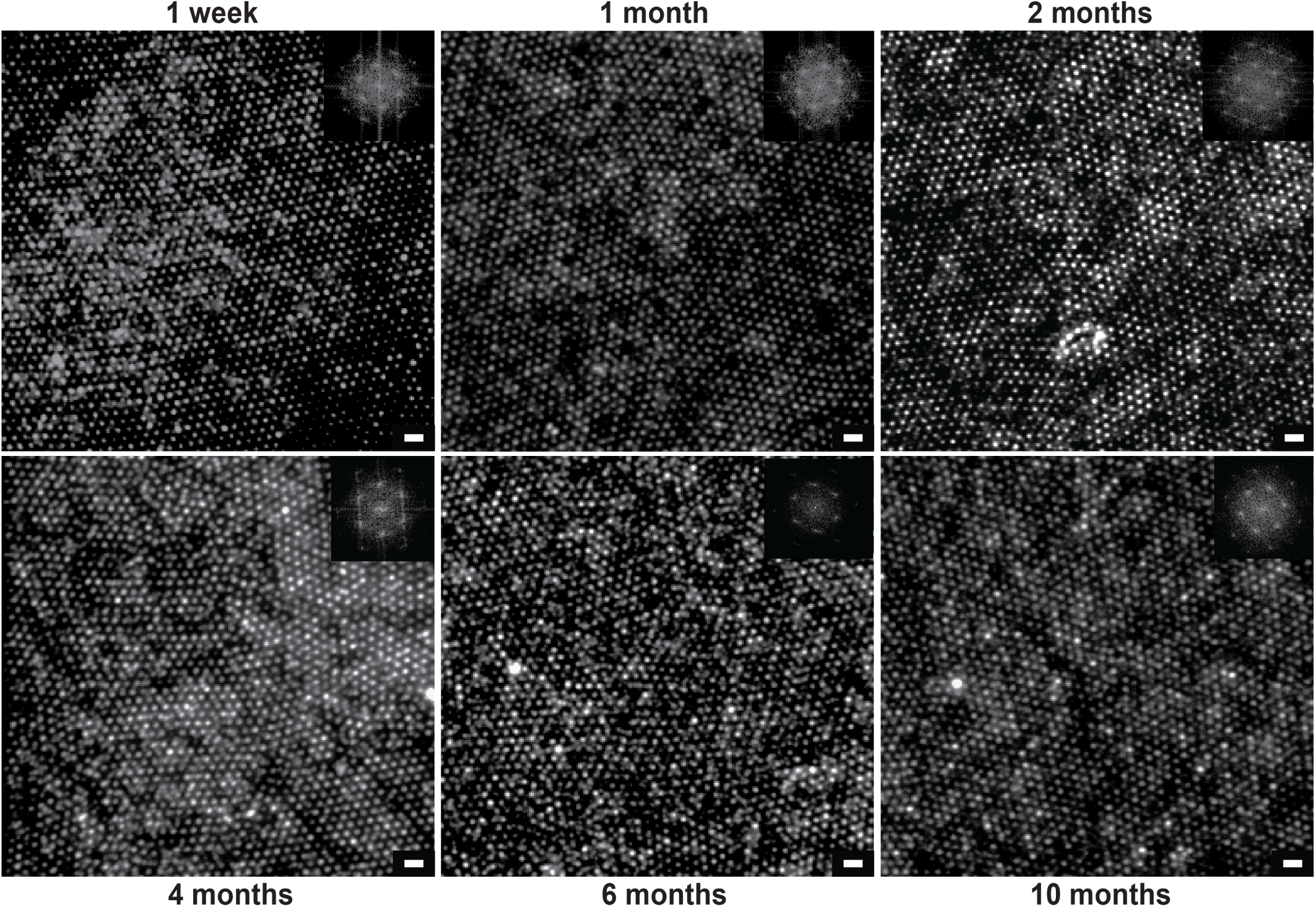
The functional viability as well as the quality of stored DNA origami nanoarrays over several months. Fluorescence images of DNA arrays after indicated storage time. DNA nanoarray shelf-life was validated using two chips for each time point: one labeled with fluorophores, and the other unlabeled. The chips were covered in aluminum foil and stored in a drawer at room temperature for up to 10 months. At each time point until 2-months, both chips were visualized with the second being labeled immediately prior to observation in 1x TAE, 12.5 mM Mg^2+^, 0.05% Tween–20. Post 2-months, already labeled chips were visualized every month for quality assessment. Scale bars are 1 µm.

**Fig. S7.**
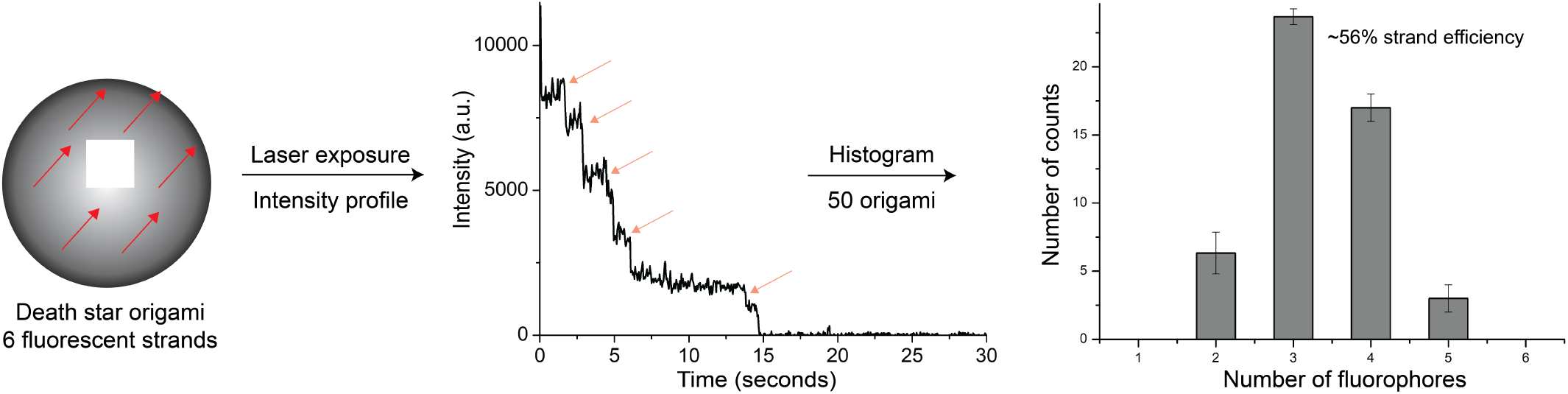
Conjugation efficiency of fluorophore-labeled strands of DNA origami nanoarray samples. (Left) Six, 20-nt sequences (same locations as PAINT docking sequences) detailed in **Table S1** hybridize complementary, fluorophore-labeled strands (≤ 10x excess) for 30 min in 1x TAE, 12.5 mM Mg^2+^, 0.05% Tween–20. (Middle) The fluorophores were photobleached over several min in the imaging buffer until all puncta disappeared. The intensity profiles of each DNA origami molecule were analyzed and the number of steps, corresponding directly to the number of strands conjugated to the origami baseplate, are counted using imageJ and iSMS. (Right) Finally, these steps are converted into a histogram to quantify strand incorporation/accessibility. For hexagonal vertices spaced 45-nm from each other, the strand conjugation efficiency is ∼56%, *i.e.* 3.36 strands of a possible 6 (*N* ≥50).

**Fig. S8.**
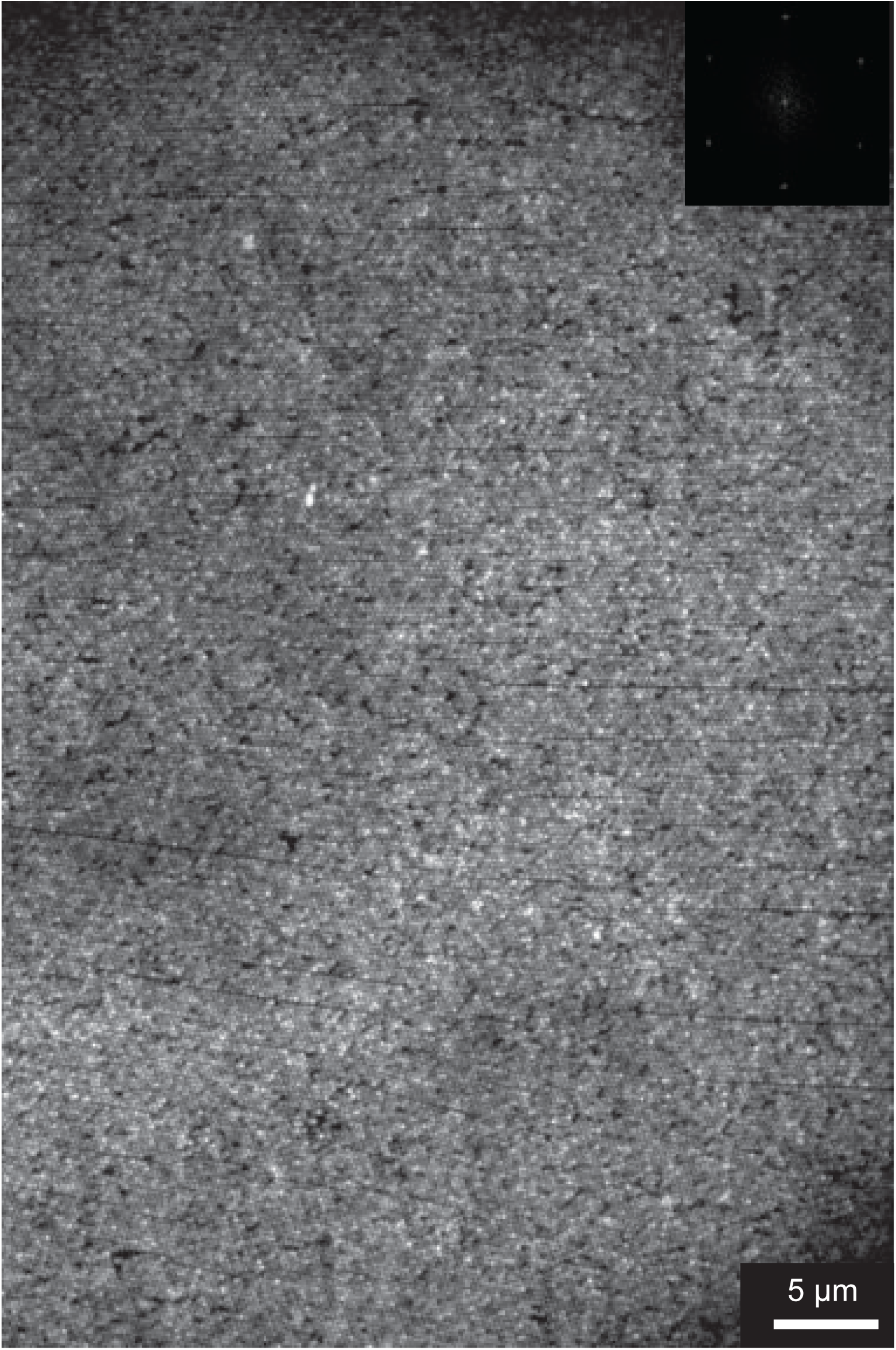
A fluorescence image of 11,000 frames collapsed along the *z*-axis of a patterned PAINT dataset prior to drift-correction and data analysis on Picasso^11^. DNA origami nanostructures were placed on a grid of binding sites created by 350 nm nanospheres. (Inset) The FFT spectrum of the image confirms the hexagonal arrangement of DNA origami.

**Fig. S9.**
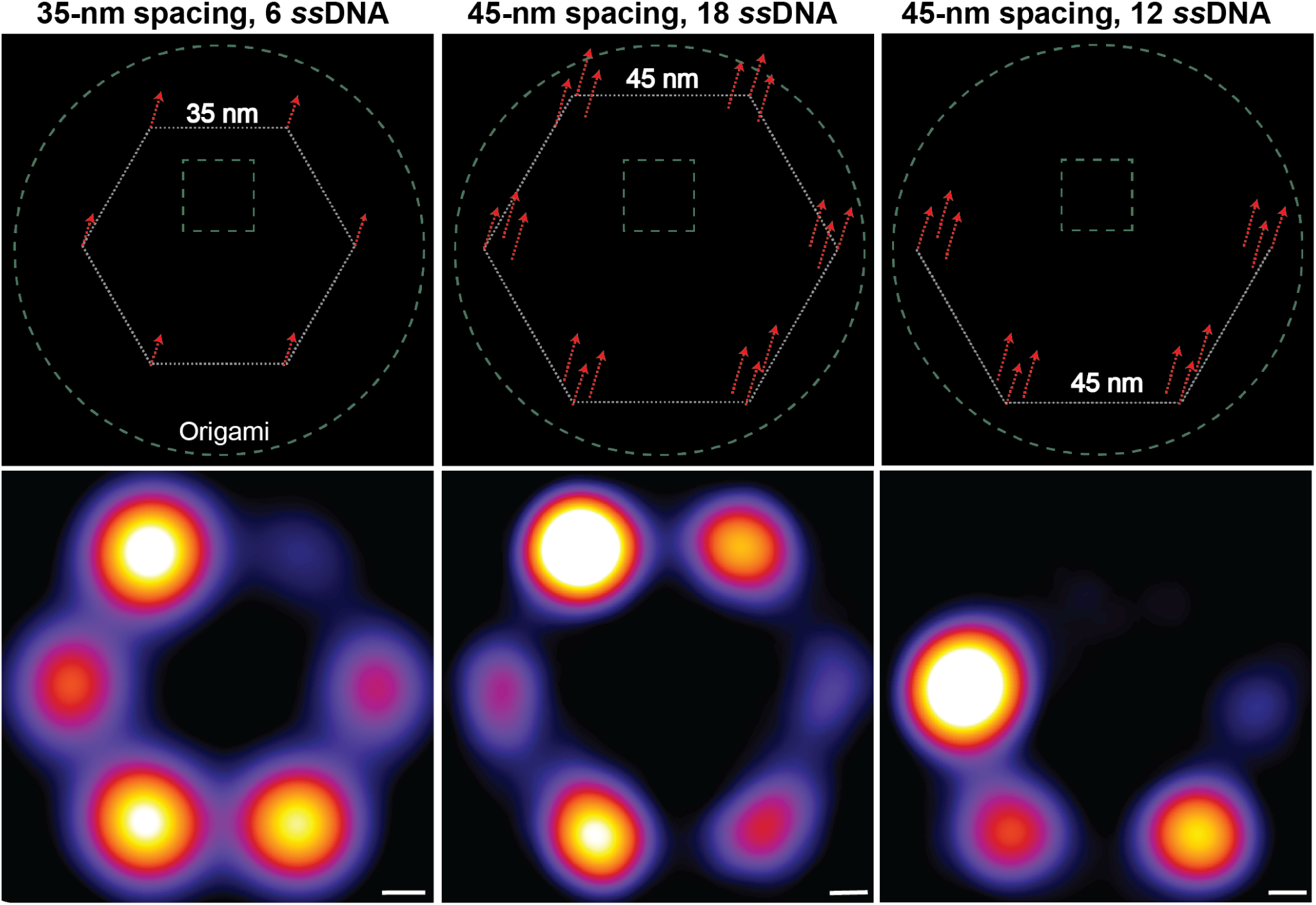
Three configurations of DNA–PAINT experiments with indicated spacings as well as number of “docking” strands (6, 18, and 12) and their corresponding averaged images formed using 10 iterations at an oversampling of 200 on Picasso^11^. From left-to-right: Manually picked structures in Picasso from a Patterned sample (*N* =300); Randomly-immobilized sample (control; *N* =200 (middle); *N* =100 (right)). Each picked structure consists of at least 4 out of 6 vertices for the 6, and 18-vertices samples, and at least 3 out of 4 vertices for the 12 vertices sample. The last two columns use redundancies of docking strands to compensate for the 56% strand conjugation efficiency (**Fig. S7**). The first column combines a larger number of structures owing to the high-density of data available in experiments with DNA origami nanoarrays.

**Fig. S10.**
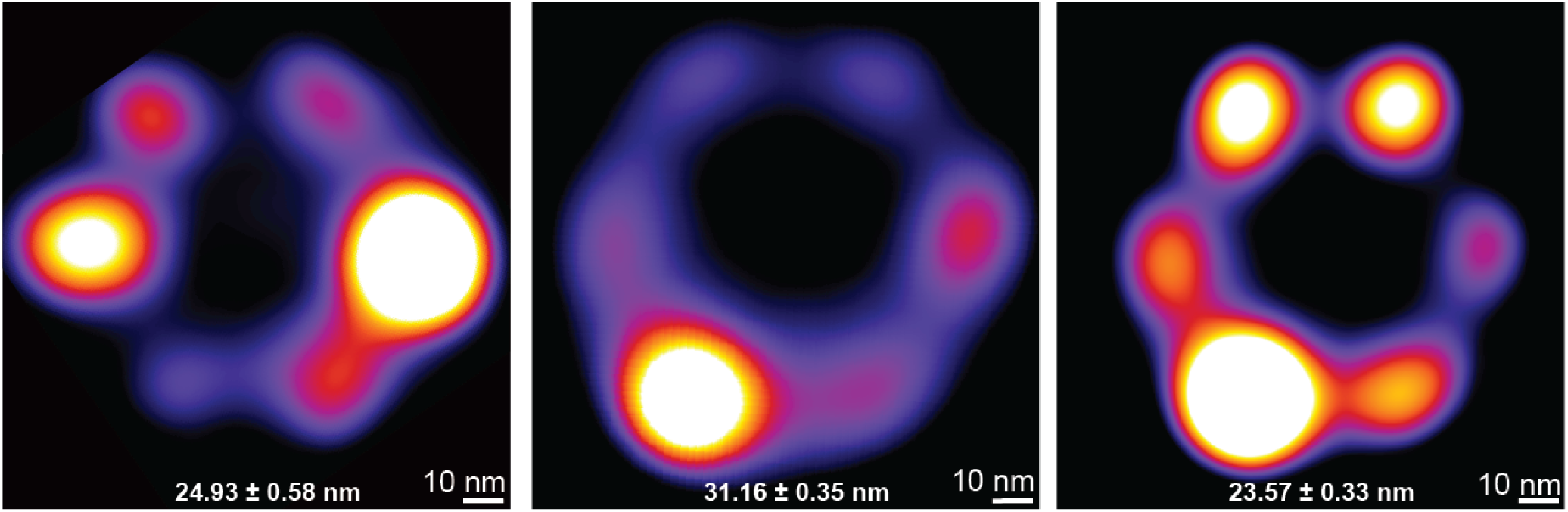
An “averaged” image of automatically-picked structures corresponding to the low, high, and patterned experimental designs (*N* = 1800, 8000, and 5200, respectively). FWHM error expressed as SEM (**Fig. 5**). The data quality in the patterned case is similar to the case with low concentration, indicating that the overlapping of multiple structures in the high concentration case might lead to “false positive” particle picking and averaging by the automated program process flow in Picasso^11^. Notably, the theoretical improvement in throughput from the low concentration case to the patterned case is ≥10x.

### S10. Supplementary Tables

**Table S2.**
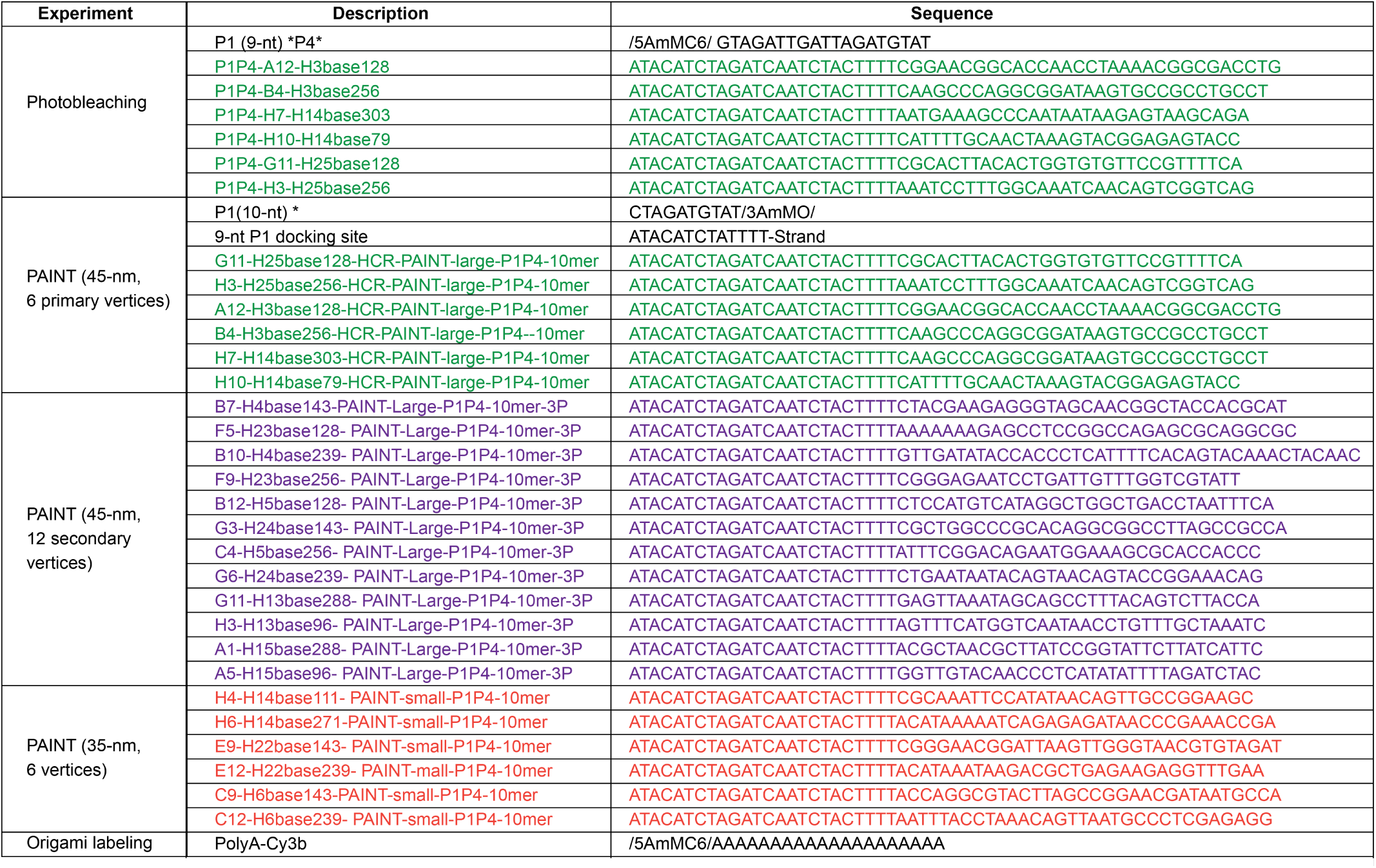
Modified strands for single-molecule experiments. The computer-aided design file, list of staples, and the staple map are included as a zip archive: Origami designs+staples+movie.zip.

**Table S3.**
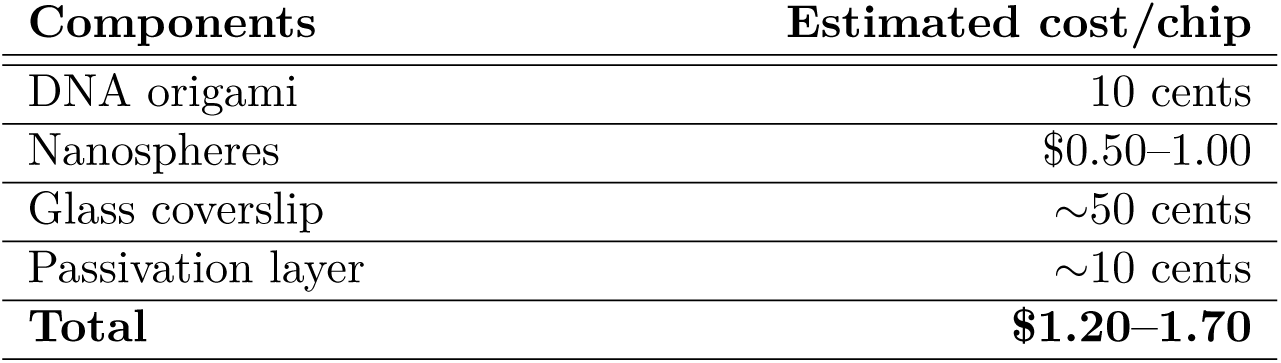
Conservative estimate of the cost of nanosphere lithography-based origami patterning materials and reagents.

| Components | Estimated cost/chip |
| --- | --- |
| DNA origami | 10 cents |
| Nanospheres | \$0.50–1.00 |
| Glass coverslip | ~50 cents |
| Passivation layer | ~10 cents |
| **Total** | **\$1.20–1.70** |

**Table S4.**
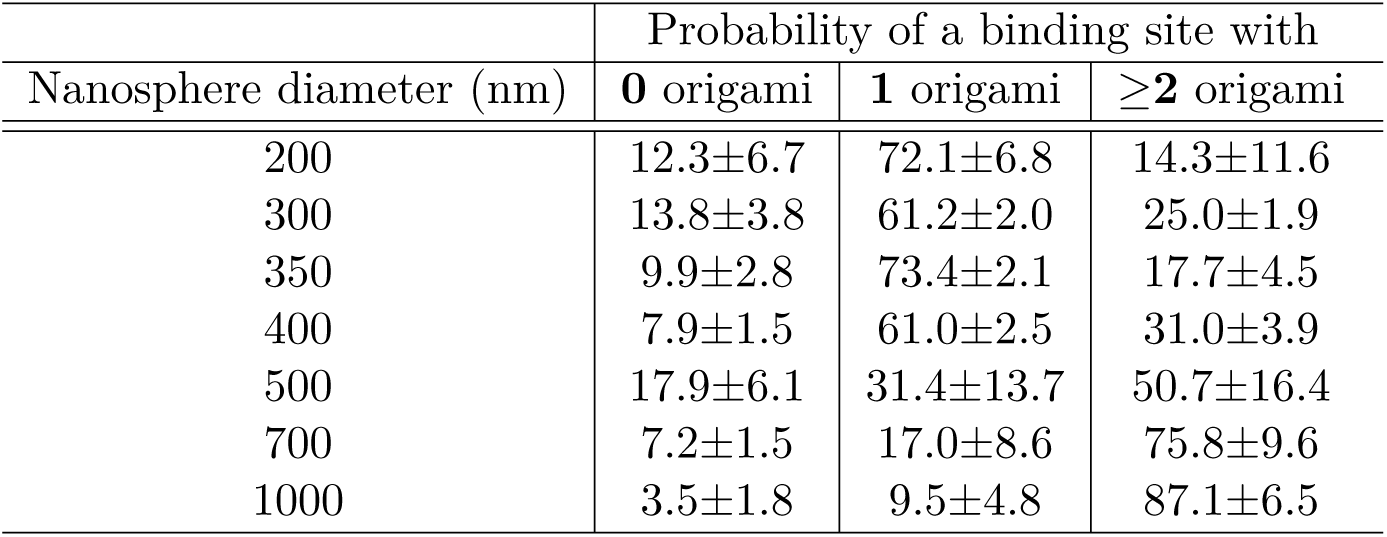
Origami binding statistics (mean±SD %) for nanosphere diameters of 200–1000 nm in **Fig. 3O**.

### S11. Supplementary Movie

Movie S1. Raw DNA–PAINT data at 100 fps showing transient, stochastic binding of 5 nM P1 imager strands with DNA origami patterned at a 350-nm pitch (300 ms exposure, 13,000 frames). Refer to zip archive: `Origami designs+staples+movie.zip` for a .avi file.

